# Pheromone-based animal communication influences the production of somatic extracellular vesicles

**DOI:** 10.1101/2022.12.22.521669

**Authors:** Agata Szczepańska, Katarzyna Olek, Klaudia Kołodziejska, Jingfang Yu, Abdulrahman Tudu Ibrahim, Laura Adamkiewicz, Frank C. Schroeder, Wojciech Pokrzywa, Michał Turek

## Abstract

Extracellular vesicles (EVs) are involved in multiple biological processes; however, there is limited knowledge of the influence of environmental factors or other individuals in a population on EV-regulated systems. The largest evolutionarily conserved EVs, exophers, are a component of the *C. elegans* maternal somatic tissue resource management system induced by embryos developing *in utero*, and thus progeny of individuals with active exopher biogenesis (exophergenesis) appear to be privileged. Using this model, we investigated the inter-tissue and social regulatory mechanisms of exophergenesis. We found that the predominant male pheromone, ascr#10, increases exopher production in hermaphrodites via the G-protein-coupled receptor STR-173 in the ASK sensory neurons. In contrast, pheromones from other hermaphrodites in the population temper exophergenesis. Within the hermaphrodite, an increase in embryo accumulation drives pro-exopher signals, and the internal sensory neurons AQR, PQR, and URX play a central role in modulating exopher levels. This intricate process is regulated partly via the neuropeptides FLP-8 and FLP-21, which originate from the URX and AQR/PQR/URX neurons, respectively. Our results reveal a regulatory network integrating internal and external cues, including control of somatic EVs production by the nervous system in response to social signals.

## Introduction

Extracellular vesicles (EVs) are lipid-bilayer-enclosed particles released by most cell types. Two major EV types can be distinguished based on their biogenesis; endosome-derived exosomes and membrane-derived ectosomes^1^. EVs can be employed by cells to remove unwanted biological material, such as misfolded proteins and damaged organelles, or to transport cargo, including proteins and nucleic acids, enabling exchange and communication between cells. Therefore, they are critical in multiple physiological processes and pathological states involving disrupted cellular homeostasis^2–5^. A recently discovered class of the largest membrane-derived evolutionarily conserved EVs, termed exophers, mediate both the elimination of waste from cells and the exchange of biological material between tissues - they were shown to play a significant role in cellular stress response, tissue homeostasis, and organismal reproduction^6–9^. Previous studies have spotlighted the role of exophers in safeguarding neuronal activity in *C. elegans* against proteotoxic stress by expelling cellular waste into surrounding tissues^7^. This mechanism is also mirrored in mammalian contexts, where Nicolás-Ávila et al. discovered that mouse cardiomyocytes release exophers loaded with defective mitochondria, thereby preventing extracellular waste accumulation and consequent inflammations, thus upholding metabolic balance within the heart^9^. However, the biological roles of exophers extend beyond the elimination of superfluous cellular components. In our previous work, we showed that the body wall muscles (BWMs) of *C. elegans* release exophers that transport muscle-synthesized yolk proteins to support offspring development, increasing their odds of development and survival^6^. However, how exophergenesis is regulated in response to external factors impacting animal development and reproduction is unclear. *C. elegans* exhibits a wide range of social behaviors governed by chemically diverse pheromones. Interwoven with sensory neuron responses, this pheromonal communication influences key developmental factors like growth, generation time, and maternal provisioning^10–12^. Building on this premise, our present study delves into understanding how these social cues modulate exophergenesis. Our findings reveal intriguing dynamics – the male-released pheromone, including ascr#10, promotes exopher formation, whereas hermaphrodite-released pheromones, including ascr#18, do the opposite, and thus population density plays a vital role in regulating vesicle production. We determined that various sensory neurons and the G-protein coupled receptor STR-173 regulate exophergenesis in response to environmental stimuli and pheromones. We also demonstrated that the AQR/PQR/URX neurons, which are directly exposed to the body cavity and monitor the worm’s body interior, restrict muscle exopher production. Taken together, we uncovered the basis for exophergenesis regulation to tune its activity to external conditions. To our knowledge, this study is the first to report how animal communication influences somatic EVs production. This study provides a deeper understanding of *C. elegans* physiology and could illuminate physiological mechanisms preserved across various species.

## Results

### Male pheromones promote exopher generation in hermaphrodite muscles

Since muscle exophers mediate the transfer of maternal resources to offspring supporting their development, we hypothesized that exophergenesis (Fig. 1a) is regulated by metabolites-derived social cues generated within the worm population. We probed this by investigating how the presence of males could impact exopher production. To this end, we examined the number of exophers in *him-5* mutants, characterized by a marked increase in the percentage of males in the population (approximately 33%, compared to 0.3% in the wild type)^13^. Interestingly, *him-5* mutant hermaphrodites co-cultured with *him-5* mutant males until the L4 stage and then transferred to a male-free plate (Fig. 1b) produce approximately 2.5 times more exophers than wild-type hermaphrodites grown on a male-free plate (Fig. 1c). This augmentation could be mediated by the previously postulated embryo-maternal signaling^6^ as *him-5* mutant hermaphrodites contain 26% more embryos *in utero* than their wild-type counterparts (Fig. 1d). To exclude the possibility that the observed increase in exopher numbers resulted from the *him-5* mutation rather than male presence, we cultured wild-type hermaphrodites on plates conditioned with *him-5* mutants or wild-type males for 48 hours, subsequently removing them (Fig. 1e). Growing hermaphrodites on male-conditioned plates increased exopher production (Fig. 1f) to the same degree as when hermaphrodites were grown with males until the L4 larvae stage (Fig. 1c). This elevation in exophers production also coincided with a rise in the number of *in utero* embryos (Fig. 1g), indicating that *C. elegans* male-secreted compounds, e.g., pheromones, can promote embryo retention in the hermaphrodite’s uterus and increase exophergenesis. Longer exposure to male-conditioned plates or co-culture with males beyond larval development did not further increase exopher production in *him-5* mutant hermaphrodites (Supplementary Fig. 1a-c). We next evaluated whether the rise in exopher production was linked to continuous or stage-specific, acute exposure to male secretions by growing hermaphrodites on male-conditioned plates during four 24-hour intervals (Fig. 1h). Results show that 24 hours of exposure to male-conditioned plates is sufficient to increase exophergenesis in hermaphrodite, regardless of the life stage at which the exposure occurred (Fig. 1h). This implies that exposure to male secretions solely during worms’ larval development can influence exopher level in adult hermaphrodites, with the most potent effect observed in animals exposed during the L4 larval stage to young adults’ day 1 stage, mirroring the effects of continuous exposure to males’ secretions (Fig. 1h). In all experiments, except for exposure during L4 larval stage to young adults’ day 1stage, increased exopher production was observed to coincide with higher embryo retention in the uterus (Fig. 1i). Our data indicate that exophergenesis is finely modulated by perception of male-derived signals, possibly pheromones, acting through embryo-maternal signaling.

**Figure 1.**
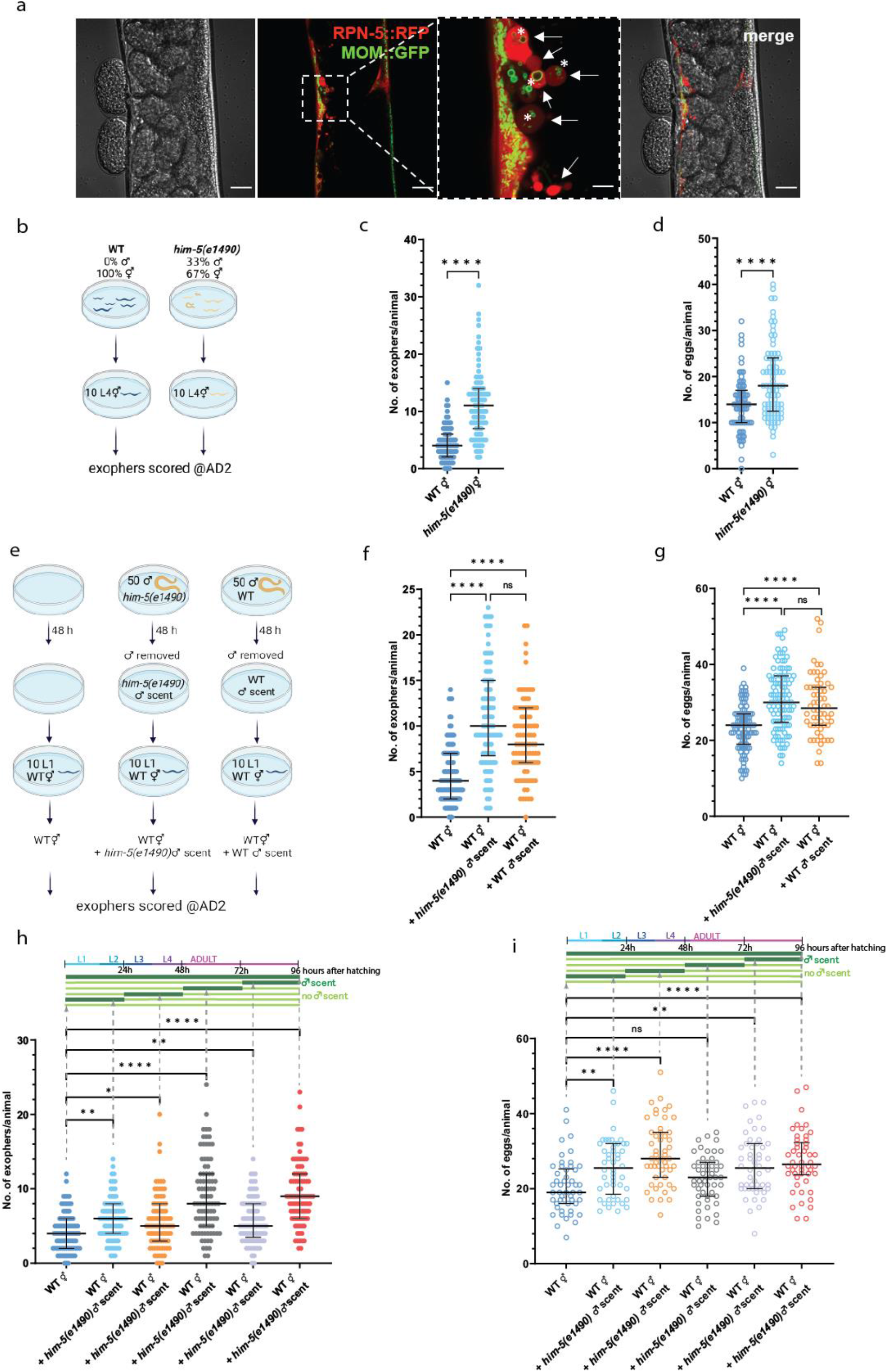
Production of *Caenorhabditis elegans* muscle exophers is regulated by male pheromones. **a** An example of *C. elegans* muscle exopher containing mitochondria. Red fluorescence comes from the RPN-5 proteasome subunit tagged with wrmScarlet fluorescent protein. Green fluorescence shows mitochondrial outer membrane (MOM) tagged with GFP. Arrows point to examples of exophers, asterisks indicate mitochondria-containing exophers. **b** Schematic representation of the experimental setup for investigating the influence of increased male presence in the population. Him-5 mutant hermaphrodites and him-5 mutant males were co-cultured from the L1 stage until the L4 stage and then transferred to a male-free plate. Hermaphrodites were grown on a male-free plate until adulthood day 2 (AD2) when the number of exophers was assessed. **c** Increased percentage of males in the population (via *him-5(e1490)* mutation) leads to a higher exophergenesis level in hermaphrodites. n = 79 and 83; N = 3 **d** Increased percentage of males in the population (via *him-5(e1490)* mutation) causes embryo retention in hermaphrodites. n = 79 and 85; N = 3 **e** Schematic representation of the experimental setup for investigating the influence of male’s secretome on exophergenesis level. 50 *him-5* or wild-type males were grown on a plate for 48 hours before being removed. Next, 10 hermaphrodites were transferred to plates previously occupied by males and grown until adulthood day 2 (AD2), when the number of exophers was assessed. **f** Growing wild-type hermaphrodites on *him-5(e1490)* or wild-type male-conditioned plates increases exophergenesis levels. n = 89 - 106; N = 3. **g** Exposing wild-type hermaphrodites to *him-5(e1490)* or wild-type male secretome causes embryo retention in the uterus. n = 60 - 105; N = 3. **h - i** Exposure to male pheromones for just 24 hours triggers exophergenesis in hermaphrodites, regardless of the life stage at which the exposure occurs. Higher embryo retention in the uterus coincided with an increase in exopher production. (h) n = 85 - 90; N = 3, (i) n = 48 – 50; N = 2. Data information: Scale bars are 20 μm and 5 μm (zoom in). @AD2 means “at adulthood day 2”. Data are presented as median with interquartile range; n represents the number of worms; N represents a number of experimental repeats that were combined into a single value; ns - not significant (P > 0.05), * P < 0.05, ** P < 0.01 P < 0.0001; (c, d) Mann-Whitney test, (f - i) Kruskal-Wallis test with Dunn’s multiple comparisons test.

### Elevated population density adversely affects exopher production

In light of our previous findings demonstrating an increase in exopher production in the presence of males within the population, we sought to examine the effect of hermaphrodite population density on this phenomenon. To this end, we established two distinct experimental conditions. In the first condition, hermaphrodites were cultured individually on plates, eliminating potential social cues. In the second condition, worms were raised as groups of 10- hermaphrodites per plate from the beginning of their development (Fig. 2a). In contrast to the effects of the presence of males, our results reveal that hermaphrodites grown at 10 hermaphrodites per plate consistently released fewer exophers compared to those grown as solitary animals (Fig. 2b). Both experimental groups contained an equivalent number of embryos *in utero* (Fig. 2c), suggesting that this modulation of exophergenesis occurs independently of embryo-maternal signaling^6^. Furthermore, we observed that cultivating hermaphrodites in a population as small as five worms per plate was sufficient to decrease exopher production (Fig. 2d). Conversely, escalating the population size to one hundred animals per plate did not cause any further significant reduction in exopher production relative to 10 hermaphrodites per plate (Fig. 2e). Growing a single hermaphrodite on a plate conditioned by other hermaphrodites reduced exopher numbers to a level comparable to that observed at a density of 10 hermaphrodites per plate (Fig. 2f). Furthermore, raising adult hermaphrodites alongside developing larval hermaphrodites significantly decrease exopher production (Fig. 2g), highlighting the importance of population density, as even the presence of immature hermaphrodites can influence exophergenesis. In summary, these findings illuminate a complex interplay between social cues and exopher production in *C. elegans*, with the presence of hermaphrodites leading to a reduction in exopher release. The effect of hermaphrodites contrasts with the observed male-induced increase, highlighting a differential response to social cues from different sexes within the population.

**Figure 2.**
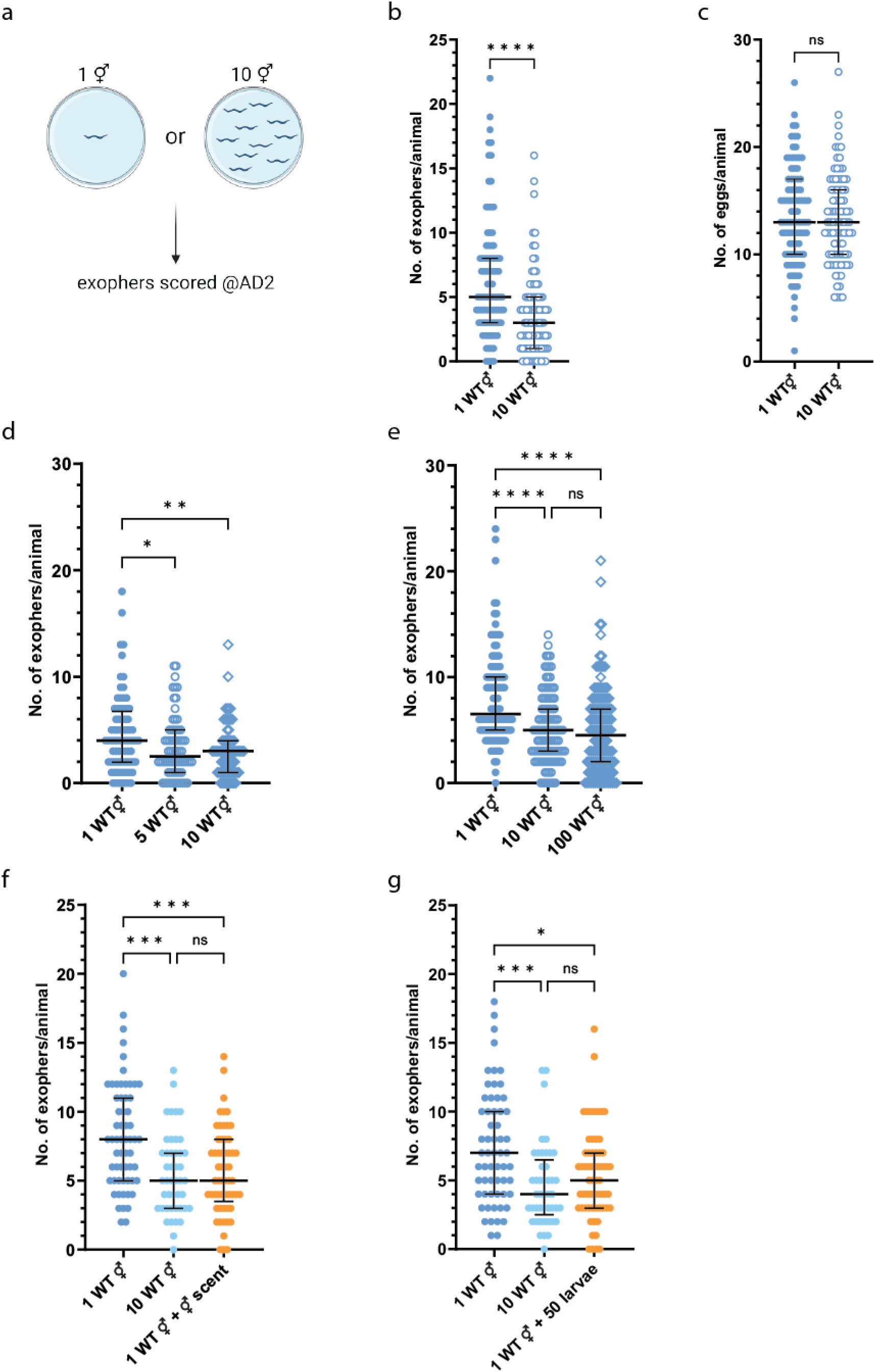
A rise in the population density of hermaphrodites adversely affects exophergenesis. **a** Experimental design for determining the effect of the presence of other hermaphrodites on the number of exophers. **b** *C. elegans* hermaphrodites grown in the presence of other hermaphrodites produce fewer exophers than solitary animals. n = 88 and 90; N = 3. **c** Embryo-maternal signaling is not responsible for the differences in exophergenesis levels as solitary and accompanied hermaphrodites contain the same number of embryos *in utero*. n = 88 and 90; N = 3. **d – e** Cultivating as few as five hermaphrodites per plate reduced exopher production, with no significant additional reduction observed when increasing the population to one hundred per plate. (d) n = 70 – 76, (e) n= 100 - 124; N = 3 **f** A single hermaphrodite on a plate previously occupied by other hermaphrodites yielded exopher numbers comparable to worms in the ten-hermaphrodite population. n = 45 and 60; N = 3 - 4. **g** Growing single adult hermaphrodite with developing larval hermaphrodites leads to a significant reduction in exopher production. n = 45 and 61; N = 3 - 4. Data information: Data are presented as median with interquartile range; n represents the number of worms; N represents a number of experimental repeats that were combined into a single value; ns - not significant (P > 0.05), * P < 0.05, ** P < 0.01, *** P < 0.001, **** P < 0.0001; (b, c) Mann-Whitney test, (d - g) Kruskal-Wallis test with Dunn’s multiple comparisons test.

### The male pheromone ascr#10 stimulates hermaphrodite exopher production

Based on our finding that male-conditioned plates are sufficient to increase muscle exopher production in hermaphrodites, we hypothesized that male-secreted pheromones may underlie the observed effects. Males secrete a wide range of sex-specific metabolites^14,15^, including several members of the ascaroside family of pheromones, among which ascr#10 is the most abundant^16^. To investigate the hypothesis that muscle exopher production in hermaphrodites may be regulated by male-derived ascaroside pheromones, we took advantage of several worm mutants in which the biosynthesis of the ascaroside side chains via peroxisomal β-oxidation is perturbed^17^. As shown in Figure 3a *acox-1(ok2257)*, *maoc-1(ok2645)*, and *daf-22(ok693)* mutants each exhibit distinct ascaroside profiles, including dramatically different amounts of the male pheromone ascr#10^18^. Similar to the male co-culture experiments, we examined exophergenesis in wild-type animals co-cultured with ascaroside biosynthesis mutants (one wild-type worm with nine mutant worms per plate, Fig. 3b). We found that co-culture with *acox-1(ok2257)* animals boosted wild-type exopher levels, whereas *maoc-1(ok2645)* reduced levels, and *daf-22(ok693)* had no significant effect (Fig. 3c), with embryo counts *in utero* remaining unaffected by all three strains (Fig. 3d), suggesting that regulation of exopher production by hermaphrodite-released pheromones is largely independent of embryo-maternal signaling.

**Figure 3.**
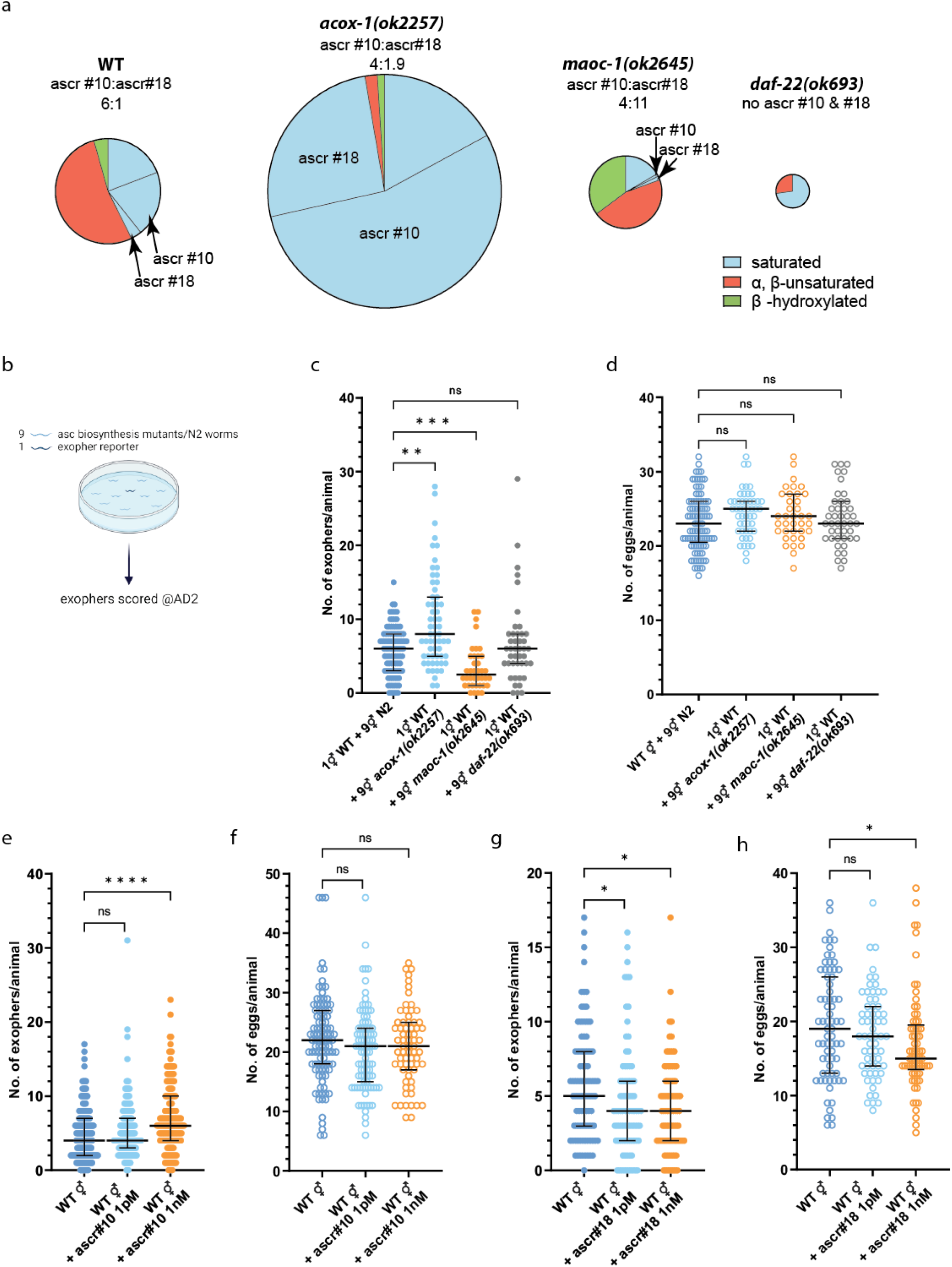
The production of exopher in hermaphrodite is stimulated by ascr#10 and inhibited by ascr#18 pheromones. **a** Mutants with altered very long chain ascarosides biosynthesis exhibit unique ascaroside pheromone profiles, characterized by varying levels of ascr#10 and its ratio to ascr#18. Based on the data from ^18^. Circle diameter denotes the proportion of produced ascarosides relative to wild-type. **b** Schematic representation of the experimental setup for investigating the influence of ascaroside side-chain biosynthesis mutants presence on exophergenesis level in wild-type worms. **c** Growing wild-type hermaphrodites in the presence of *acox-1(ok2257)* and *maoc-1(ok2645)* mutant increase and decrease exopher production, respectively. n = 38 - 93; N = 3. **d** Growing wild-type hermaphrodites with ascaroside side-chain biosynthesis mutants does not influence the number of embryos in the uterus. n = 37 - 93; N = 3. **e** Hermaphrodites grown on NGM plates supplemented with 1 ng of ascr#10 have increased levels of exophergenesis. n = 164 - 190; N = 5 - 6. **f** ascr#10 do not influence embryo retention in hermaphrodites. n = 69 - 101; N = 4. **g** Hermaphrodites grown on NGM plates supplemented with 1 pg or 1 ng of ascr#18 have decreased levels of exophergenesis. n = 83 - 90; N = 3. **h** ascr#18 do not influence embryo retention in hermaphrodites. n = 63 - 71; N = 3. Data information: @AD2 means “at adulthood day 2”. Data are presented as median with interquartile range; n represents the number of worms; N represents a number of experimental repeats that were combined into a single value; ns - not significant (P > 0.05), * P < 0.05, ** P < 0.01, *** P < 0.001, **** P < 0.0001; Kruskal-Wallis test with Dunn’s multiple comparisons test.

Observing an increased exopher count when wild-type hermaphrodites were exposed to ascarosides released by *acox-1(ok2257)* hermaphrodites, known for their elevated ascr#10 levels^18^, and considering the dominance of ascr#10 in male pheromone profiles^16^, we assessed the role of this specific ascaroside in exopher upregulation. Remarkably, we found that exposure to low concentrations of ascr#10 (∼ 1 nanomolar) led hermaphrodites to produce more exophers than control animals, while the number of embryos in the uterus remained unchanged (Fig. 3e-f). Ascr#10 alone amplifies exophergenesis, yet the *acox-1* mutants also exhibit a notable presence of ascr#18 alongside it^18^. Prompted by this, we examined ascr#18’s role in exopher modulation. Unexpectedly, exposure of animals to concentrations as low as approx. 1 picomolar of ascr#18 reduced exopher formation in reporter hermaphrodites (Fig. 3g), and at greater concentrations (∼ 1 nanomolar), it also reduced egg retention in utero (Fig. 3h). These results show that different ascaroside pheromones have starkly different effects on exopher production, emphasizing the intricate relationship between chemical structure and activity in ascarosides^17^.

### Bacterial diet has a limited influence on exopher production

To rule out the possibility that exophergenesis is substantially regulated by the molecules derived from the bacterial food source and to further explore the relationship between sensory neurons, pheromones, and bacterial diet in *C. elegans*, we tested various bacterial strains that might indirectly influence worm-to-worm communication. We directly compared the *Escherichia coli* B strain OP50 and K-12 strain HT115, which are prevalently utilized in *C. elegans* culture and RNAi silencing experiments, respectively^19,20^. Our findings revealed an increase in the number of exophers when the *E. coli* HT115 diet was applied, as opposed to the OP50 diet (Supplementary Fig. 2a). However, there was a notable elevation in the number of eggs present in utero on the second day of adulthood under the HT115 diet (Supplementary Fig. 2b). Importantly, the effect of the presence of males on the increase in exopher numbers was observed, regardless of the bacterial strain used for feeding (Supplementary Fig. 2c). The metabolic activity of the bacteria, whether alive or PFA-killed prior to culture, did not impact the results (Supplementary Fig. 2a), corroborating that exophergenesis in the worm’s muscles is consistently activated, irrespective of the bacterial type used as a food source.

### ASK, AWB, and ADL sensory neurons govern exophergenesis in response to males-social cues

Given the role of many olfactory neurons in detecting ascarosides^16^ (Fig. 4a), we examined whether genetic ablation of all ciliated sensory neurons would abolish male pheromone-regulated exopher induction. Indeed, as shown in Fig. 4b, *che-13* mutants, which do not form proper cilia and are incapable of pheromone detection^21^, produce a minimal number of exophers (Fig. 4b). To pinpoint the specific sensory neurons mediating pheromone-dependent modulation of exophergenesis, we examined its level in animals with genetically ablated sensory neurons previously identified as capable of pheromone detection^22^. We found that the elimination of ASK, ADL, or AWC suppressed exopher production comparable to that in *che-13* mutants, while ASH ablation led to a modest reduction (Fig. 4c). Notably, worms with genetically ablated neurons, which showed diminished exophergenesis, also displayed fewer eggs *in utero* (Fig. 4d). Considering the prominent effect on exophergenesis observed in genetic ablation of AWC thermo-responsive neurons^23,24^, we investigated the influence of temperature on exophergenesis. When growing worms at 15, 20, or 25 °C, we noted a temperature-related rise in exopher formation (Fig. 4e). This increase was inhibited in the absence of AWC, substantiating the temperature-dependent control of exophergenesis by these neurons (Fig. 4e). Interestingly, the introduction of male-specific, pheromone-sensing CEM neurons in hermaphrodites (via *ceh-30* gain-of-function mutation^25^) did not lead to changes in exopher level (Fig. 4c).

**Figure 4.**
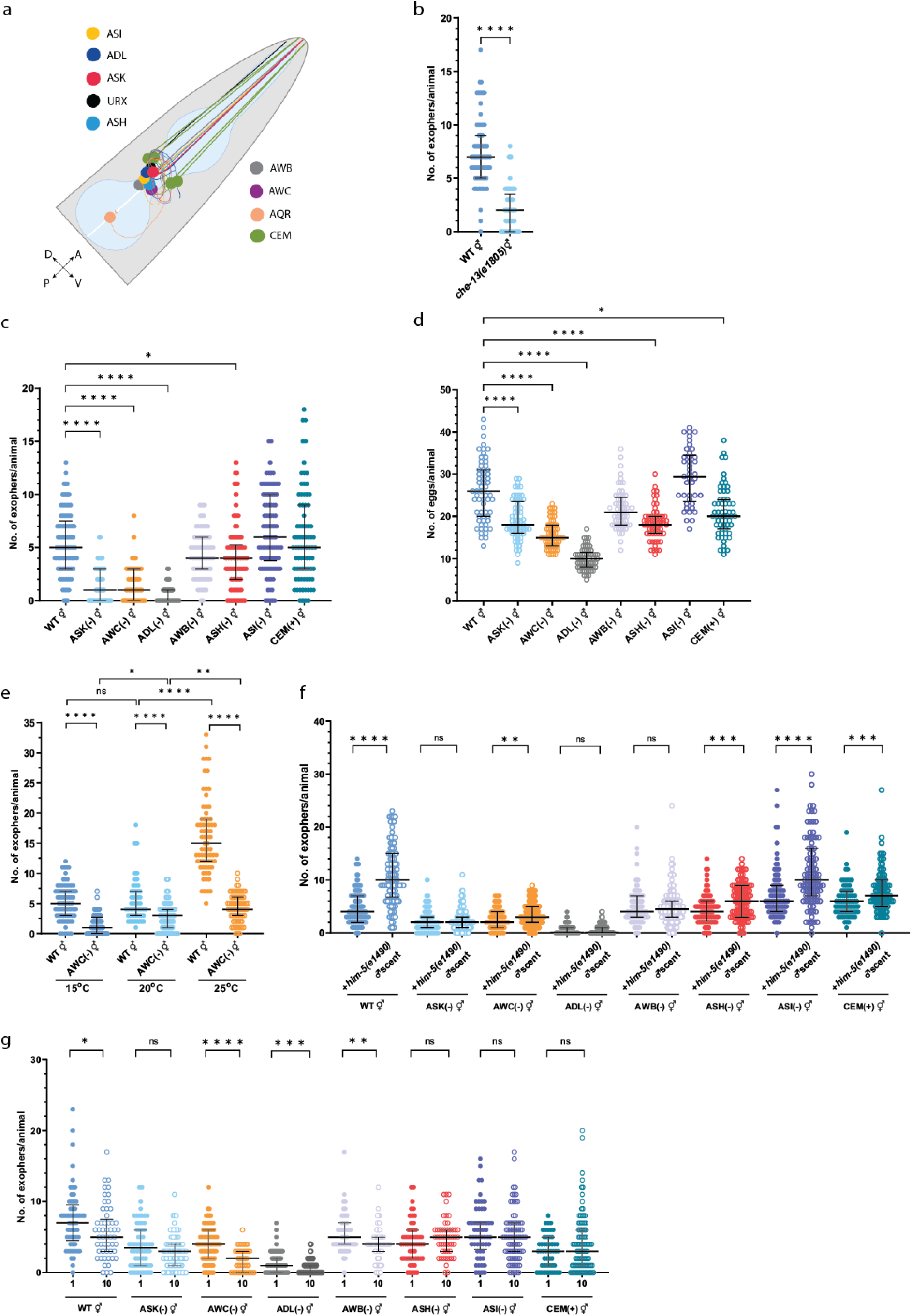
Multiple olfactory neurons regulate exophergenesis levels in response to hermaphrodite and male pheromones. **a** Sensory neurons investigated within this study. **b** Hermaphrodites with impaired ciliated sensory neurons have lower exophergenesis levels. n = 66 and 65; N = 3. **c** Genetic ablation of ASK, AWC, ADL, or ASH neurons reduces exopher production. n = 61 - 97; N = 3. **d** Worms with genetically ablated neurons, resulting in reduced exophergenesis, also exhibited fewer eggs *in utero*. n = 42 - 61; N = 2 - 3. **e** AWC neurons regulate the temperature-dependent increase in muscle exopher production. n = 52 - 92; N = 3. **f** Increase in exophergenesis levels due to exposure to male pheromones is mediated by ASK, AWB, and ADL neurons. Schematic representation of the experimental setup is presented in Fig. 1e (without conditioning with WT males). n = 77 - 108; N = 3. **g** Decreased exophergenesis level due to exposure to hermaphrodite’s pheromones is mediated by ASH, ASI, and ASK neurons and can be altered by adding CEM male-specific neurons. Schematic representation of the experimental setup is presented in Fig. 2a. n = 49 - 97; N = 3. Data information: Data are presented as median with interquartile range; n represents the number of worms; N represents a number of experimental repeats that were combined into a single value; ns - not significant (P > 0.05), * P < 0.05, ** P < 0.01, *** P < 0.001, **** P < 0.0001; (b, f, g) Mann-Whitney test, (c - e) Kruskal-Wallis test with Dunn’s multiple comparisons test.

Subsequently, we explored which olfactory neurons are essential for detecting male pheromones that enhance exophergenesis. By growing hermaphrodites with genetic ablations of different classes of olfactory neurons on *him-5* mutant male-conditioned plates (as shown in Fig. 1e scheme), we determined that the removal of ASK, AWB, or ADL neurons prevented the increase in the number of embryos *in utero* and nullified the enhancement in exopher production driven by male-emitted pheromones (Fig. 4f; Supplementary Fig. 3a). These results align with previously described roles for ASK, AWB, and ADL neurons in male pheromones sensing^26–29^. This demonstrates the critical role of the neuronal odor recognition system in modulating EV production by muscles and highlights the importance of particular neurons in recognizing distinct pheromone types.

### ASK, ASI, and ASH olfactory neurons regulate exopher production in hermaphrodite populations

We sought to pinpoint the neurons crucial for downregulating exopher production in response to the presence of other hermaphrodites. We grew hermaphrodites with genetically ablated different classes of olfactory neurons either singly or in a 10 hermaphrodite population (Fig. 2a scheme). Our analysis with strains showing impaired olfaction revealed that ASH, ASI, and ASK neurons are pivotal in this process (Fig. 4g). Remarkably, the masculinization of a hermaphrodite olfactory circuit through the introduction of CEM male-specific neurons resulted in the absence of exopher production decrease in ten hermaphrodite population comparing to solitary worms (Fig. 4g). There was no correlation between exopher numbers and eggs *in utero* count neither for wild-type animals nor for ablation mutants with (ASI) or without (AWB) impaired hermaphrodite pheromones recognition (Supplementary Fig. 3b). This further demonstrates that hermaphrodite pheromones regulate exophergenesis by omitting embryo-maternal signaling. Finally, the complex interaction between ascaroside synthesis and neural mechanisms in the regulation of exophergenesis was reinforced by the interplay observed between ASI and AWB neuronal mutant hermaphrodites with ascaroside mutant hermaphrodites (Supplementary Fig. 3c-d). In summary, the critical role of ASH, ASI, and ASK neurons underlines the nuanced regulation of exophergenesis in response to the presence of other hermaphrodites.

### STR-173 receptor mediates pheromone-dependent exophergenesis modulation through ascr#10

Binding signaling molecules to the relevant receptor is the first step in transducing neuron chemosensory signals. More than 1,300 G protein-coupled receptors (GPCRs) mediate this communication in *C. elegans*^30^. Internal states and environmental conditions can modulate GPCRs expression to affect worm behavior^31–34^. To identify the receptor(s) responsible for ascaroside-mediated alterations in exopher formation, we performed RNA-sequencing of animals grown either as a single animal or in a 10-hermaphrodites population (Supplementary Fig. 4a). On the transcriptional level, neither group differed markedly (Supplementary Fig. 4b), and none of the 7TM receptors was significant up- or down-regulated (Supplementary Table 1). However, *str-173* receptor transcript, which shows among all of the G protein-coupled receptors one of the most evident trends between growth conditions (2.3 fold change), is, according to single-cell RNA-seq data^35^, expressed almost exclusively in ASK neurons (Supplementary Fig. 4c). Since ASK is crucial for pheromone-mediated modulation of exopher formation, we investigated the *str-173* role in this pathway.

The wrmScarlet CRISPR/Cas9-mediated transcriptional knock-in for the *str-173* gene confirmed its strong expression during the late development and adulthood in ASK neurons and revealed additional expression in OLQ neurons, the pharynx, vulva muscles, and the tail (Fig. 5a-b). Next, we created *str-173* null mutants again using CRISPR/Cas9 editing (Supplementary Fig. 4d). We observed that *str-173* gene knockout neither influence the basal level of exophergenesis in *str-173* mutant hermaphrodites (Fig. 5c) nor robustly impact population density-mediated exophergenesis decrease (Fig. 5d). Interestingly, contrary to wild-type worms, *str-173* mutants did not exhibit an increase in exopher production in response to male pheromones (Fig. 5e). Recognizing the potential function of STR-173 in male pheromone-mediated signaling, we investigated its participation in the ascr#10 recognition.

**Figure 5.**
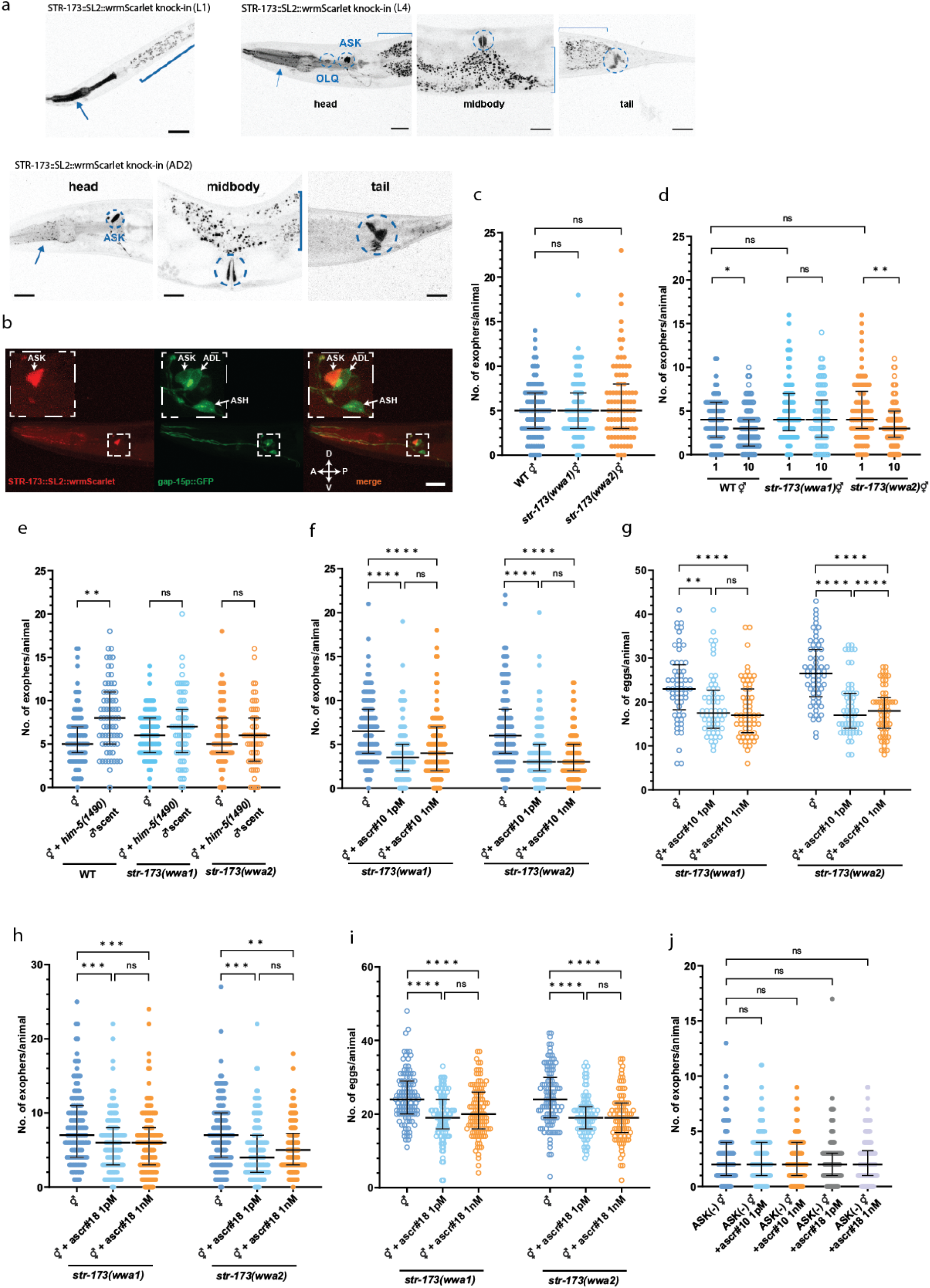
The STR-173 receptor facilitates the pheromone-dependent modulation of exophergenesis via ascr#10. **a - b** *str-173* 7TM receptor is expressed in neurons (ASK and probably^#^ OLQ) and non-neuronal tissues (pharynx marked with an arrow, vulva, and probably^##^ rectal gland marked with circles). Square brackets mark gut autofluorescence. ^#^based on the position and scRNAseq data^35^ ^##^based on the position and the shape. **c** Exophergenesis level in *str-173* mutants co-cultured with other hermaphrodites is unchanged compared to wild-type worms. n = 90; N = 6. **d** Knocking out the *str-173* gene does not roboustly affect the decrease in exophergenesis mediated by changes in population density. n = 86 - 90; N = 3. **e** Increase in exophergenesis levels due to exposure to male pheromones depends on STR-173 7TM receptor. n = 60 - 90; N = 2 - 3. **f - g** When exposed to ascr#10, *str-173* mutants exhibit a reduction in exopher production and *in utero* embryo number, underscoring the importance of STR-173 in binding ascr#10. (f) n = 104 – 113, N = 4; (g) n = 60, N = 2 **h - i** *Str-173* mutants, when cultured on NGM plates supplemented with ascr#18, experience a decrease in exopher production. (h) n = 124 - 134, N = 3; (i) n = 96 - 115, N = 3 **j** Removal of ASK neurons nullifies ascr#10 and ascr#18 influence on exophergenesis. n = 99 - 135; N = 3. Data information: Scale bars are (a) 30 μm and (b) 20 μm. Data are presented as median with interquartile range; n represents the number of worms; N represents a number of experimental repeats that were combined into a single value; ns - not significant (P > 0.05), * P<0.05, ** P < 0.01, *** P < 0.001, **** P < 0.0001; (c, d - j) Kruskal-Wallis test with Dunn’s multiple comparisons test, (d, e) Mann-Whitney test.

Contrary to the increase in exopher production observed in wild-type animals, *str-173* mutants exposed to ascr#10 display a decrease in exopher production, emphasizing STR-173’s role in translating male pheromone cues into physiological responses (Fig. 5f-g). Furthermore, *str-173* mutants grown on NGM plates supplemented with ascr#18, like wild-type worms, show a decrease in exopher production (Fig. 5h-i). This emphasizes the specificity of STR-173 binding to ascr#10 rather than any other type of ascaroside.

The observed decrease in exopher production in *str-173* mutants exposed to ascr#10 suggests the presence of another ascaroside receptor(s) capable of binding ascr#10 that plays an opposite role in exopher regulation to STR-173. Exposing worms with ablated ASK neurons to ascr#10 or ascr#18 did not lead to significant changes in exopher counts (Fig. 5j), suggesting that receptor(s) for ascr#18 and ascr#10, which decrease exopher production, are located in ASK neurons. In summary, our data indicate a vital role of ASK in exopher regulation, mediated through various ascaroside receptors expressed in ASK neurons.

### AQR, PQR, and URX neurons activity limit exophergenesis

Among the 118 classes of neurons in *C. elegans*, only four are directly exposed to the pseudocoelomic cavity^36^. Three classes of these neurons, AQR, PQR, and URX, regulate social feeding in worms^37^. Given that exophers are released to the worm’s pseudocoelomic cavity and are regulated by social cues, we hypothesized that AQR, PQR, and URX might play a role in exophergenesis. To test for that, we first investigated the effect of genetic ablation of AQR, PQR, and URX neurons on exopher production. Indeed, the removal of these neurons leads to a substantial increase in exophergenesis (Fig. 6a). Notably, the increased number of exophers generated by worms with genetically ablated AQR, PQR, and URX neurons is not the result of embryo-maternal signaling as these animals have a modestly reduced number of eggs *in utero* (Fig. 6b) and smaller brood size (Supplementary Fig. 5a-b) than wild-type control.

**Figure 6.**
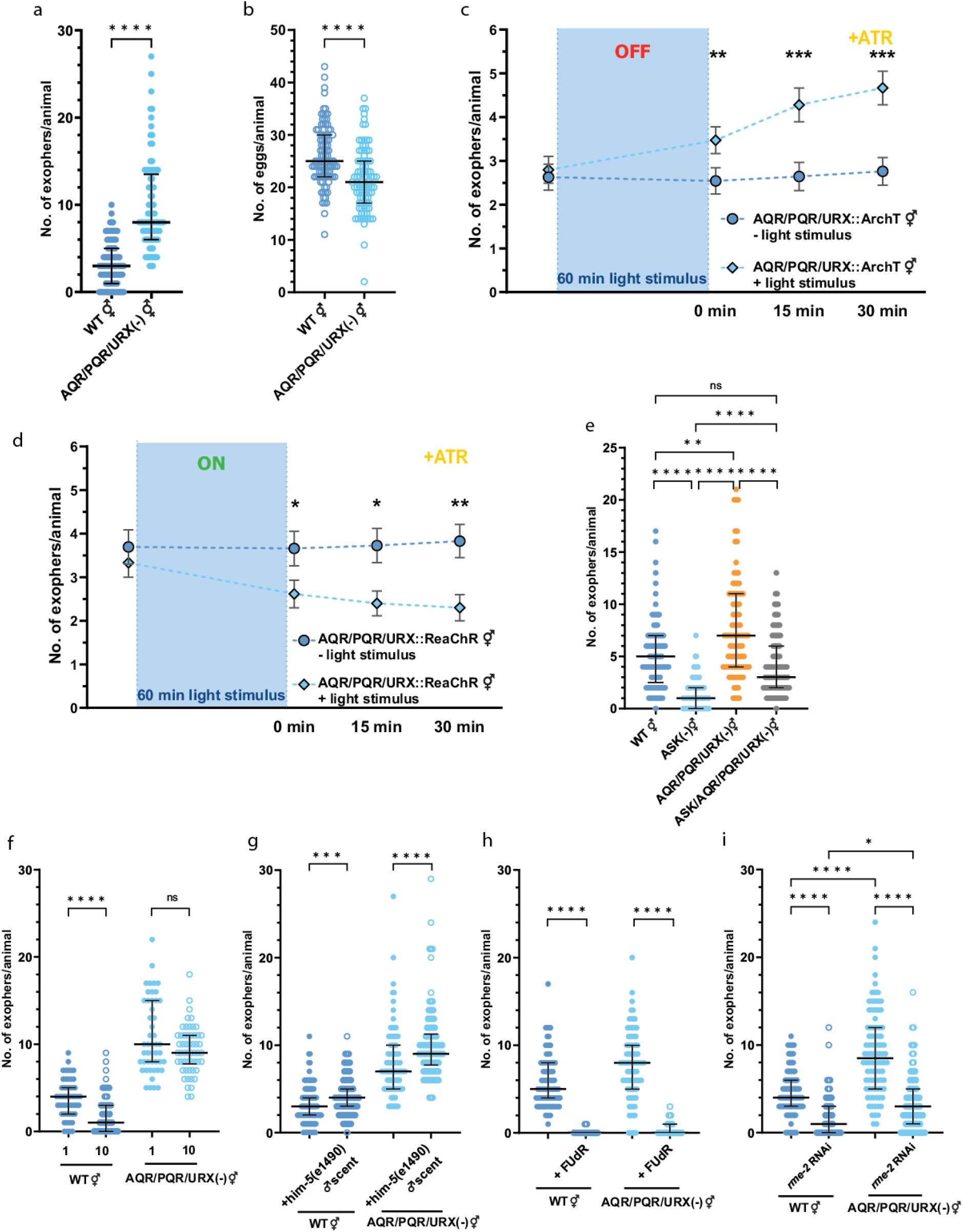
Pseudocoelom-exposed neurons negatively regulate exopher production. **a** Genetic ablation of pseudocoelom-exposed AQR, PQR, and URX neurons increases exopher production. n = 89 and 100; N = 3. **b** Genetic ablation of pseudocoelom-exposed AQR, PQR, and URX neurons causes a decrease in the number of embryos present in the uterus. n = 89 and 100; N = 3. **c** ArchT-mediated optogenetic inactivation of AQR, PQR, and URX neurons increases exopher production. n = 59; N = 6. **d** ReaChR-mediated optogenetic activation of AQR, PQR, and URX neurons decreases exopher production. n = 59 and 60; N = 6. **e** The opposing exophergenesis levels observed in animals with genetic ablation of ASK neurons (low exophergenesis) and AQR, PQR, and URX neurons (high exophergenesis) converge to an intermediate level in animals with all four neurons removed. n = 89 - 95; N =3. **f** Decreased exophergenesis level due to exposure to hermaphrodite’s pheromones is partially mediated by AQR, PQR, and/or URX neurons. n = 51 - 60; N = 3. **g** Increase in exophergenesis levels due to exposure to male pheromones is not altered in animals with genetic ablation of AQR, PQR, and URX neurons. n = 102 - 105; N = 3. **h** Genetic ablation of AQR, PQR, and URX neurons does not rescue the inhibition of exophergenesis caused by FUdR-mediated worm sterility. n = 89 - 90; N = 3. **i** Genetic ablation of AQR, PQR, and URX neurons only partially rescues the inhibition of exophergenesis caused by *rme-2* (yolk receptor) knockdown. n = 90; N = 3. Data information: +ATR means “with all-trans-retinal”. Data are presented as median with interquartile range (a – b, e – i) or mean with SEM (c – d); n represents the number of worms; N represents a number of experimental repeats that were combined into a single value; ns - not significant (P > 0.05), * P < 0.05, ** P < 0.01,*** P < 0.001, **** P < 0.0001; (a - d, f - h) Mann-Whitney test, (e, i) Kruskal-Wallis test with Dunn’s multiple comparisons test.

To further validate the role of AQR, PQR, and URX neurons in the regulation of exophergenesis, we optogenetically inactivated or activated them using ArchT^38^ or ReaChR^39^, respectively, and compared the number of exophers before and after the stimulus. We observed that 60 min of AQR, PQR, and URX neurons inactivation leads to a significant increase in exopher release (Fig. 6c and Supplementary Fig. 5c). On the other hand, 60 min of AQR, PQR, and URX neurons activation resulted in a significant decrease in exopher release after the stimulus was completed (Fig. 6d and Supplementary Fig. 5d). This modulation was evident at the adult day 1 (AD1) stage for ReaChr-based activation but not ArchT-based inactivation, underscoring the potential significance of an optimal embryo count or direct uterus interaction (Supplementary Fig. 5e-h).

Moreover, our data underscores that the contrasting exophergenesis characteristics observed in ASK-ablated worms and in worms with genetic elimination of AQR, PQR, and URX neurons resemble those of wild-type worms in animals lacking all four neuron classes (Fig. 6e). We further determined that AQR, PQR, and URX neurons are involved in response to hermaphrodite pheromones (Fig. 6f), but not to male pheromones (Fig. 6g). However, their heightened activity could not override the indispensable role of fertility in exophergenesis (Fig. 6h-i). In summary, AQR, PQR, and URX neurons distinctly modulate exopher production in response to hermaphrodite pheromones, underscoring the complexity of neuronally-regulated reproductive signaling.

### URX-expressed FLP-8 and AQR/PQR/URX-expressed FLP-21 neuropeptides inhibit exophergenesis

To elucidate the mechanism by which AQR/PQR/URX neurons regulate muscle exopher production, we used single-cell RNA-seq data^31^, targeting neuropeptides predominantly expressed in these neurons with a limited expression in other cells. These criteria were fulfilled by URX-expressed FLP-8, and AQR, PQR, URX-expressed FLP-21 neuropeptides (Supplementary Fig. 6 a-c). We could demonstrate that these neuropeptides negatively regulate exophers production, as evidenced by increased exophers counts in both single and double mutants (Fig. 7a), independent of embryo-maternal signaling (Fig. 7b). Moreover, FLP-8 and FLP-21 act downstream of URX and AQR/PQR/URX, respectively, as optogenetical activation of these neurons in the mutant context failed to suppress exopher release as observed in control animals (Fig. 7c-e, Supplementary Fig. 7a-c). The results acquired regarding the influence of the FLP-21 neuropeptide on the regulation of socially-driven exophergenesis align with the previously established role of FLP-21 in controlling social behavior^40^.

**Figure 7.**
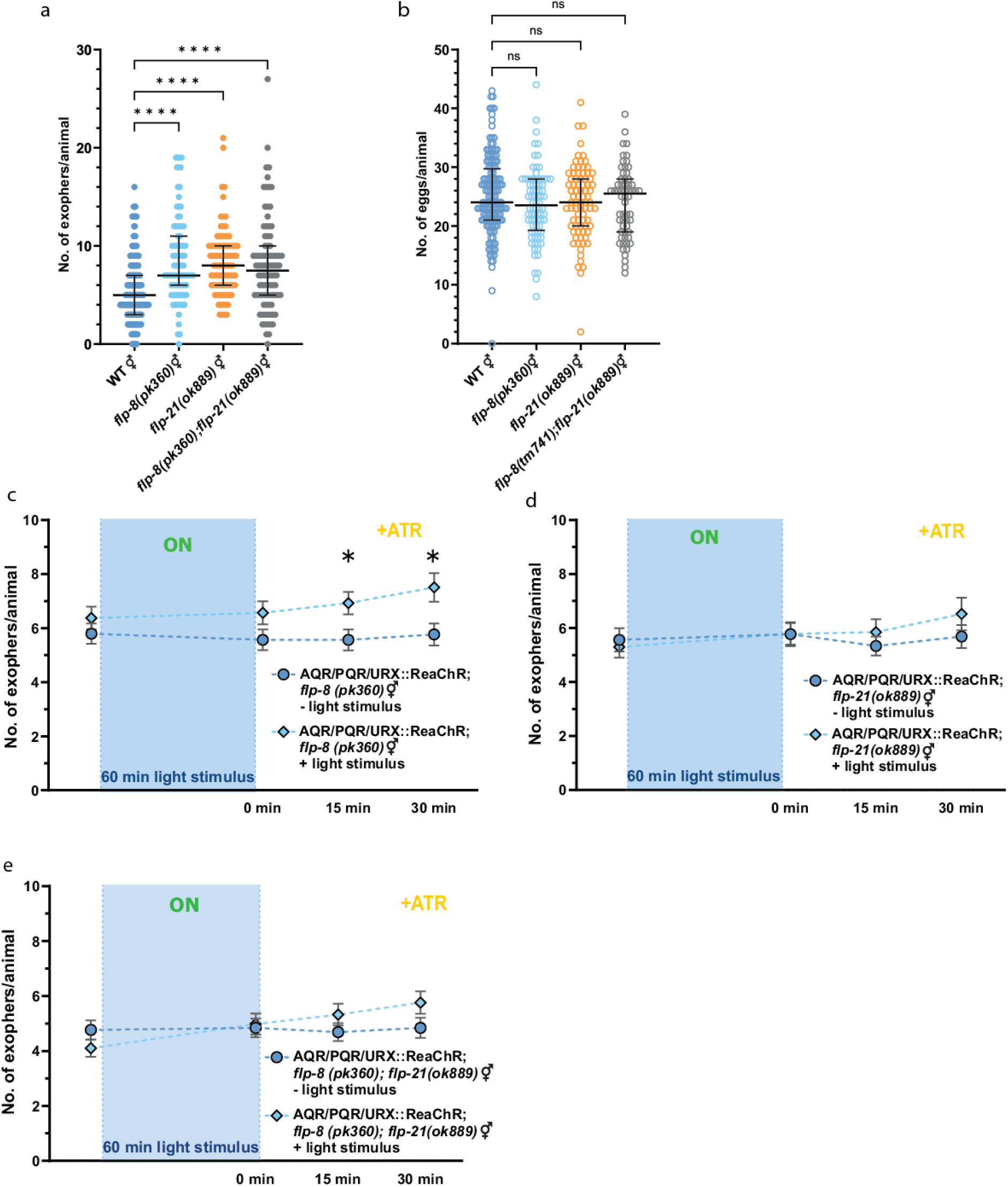
Neuropeptides FLP-8 released by URX neurons and FLP-21 released by AQR, PQR, and/or URX inhibit exophergenesis. **a** *flp-8(pk360)* and *flp-21(ok889)*single and double mutants have increased exophergenesis levels. n=180, N=6 for WT; n=90, N=3 for mutants. **b** Mutations in *flp-8(pk360)* and *flp-21(ok889)* genes do not influence embryo retention *in utero*. n=140, N=5 for WT; n=60-80, N=3 for mutants. **c-e** Optogenetic activation of AQR, PQR, and URX which do not produce FLP-8 and FLP-21 neuropeptides do not lead to a decrease in exopher production. n = 65 - 71; N = 6. Data information: +ATR means “with all-trans-retinal”. Data are presented as median with interquartile range (a – b) or mean with SEM (c – e); n represents the number of worms; N represents a number of experimental repeats that were combined into a single value; ns - not significant (P > 0.05), * P < 0.05, **** P < 0.0001; (c - e) Mann-Whitney test, (a, b) Kruskal-Wallis test with Dunn’s multiple comparisons test.

### Model of exopher regulation mechanism in *C. elegans*

Our investigations have led us to construct the following model detailing the intricate mechanism of exopher regulation:

1. Male pheromones: Males produce ascarosides, including ascr#10, that increase exopher production levels. These signals predominantly act through ASK, ADL, and AWB. The critical role in this pathway is played by ASK-expressed STR-173 G protein-coupled receptor, which binds ascr#10 to potentiate exopher production.
2. Hermaphrodite pheromones: Hermaphrodites release ascarosides, including ascr#18, that reduce exopher production levels. These signals predominantly act through ASI, ASH, and ASK (via ascr#18) sensory neurons.
3. Embryo accumulation effect: The rise in embryos inside the hermaphrodite triggers a series of pro-exopher signals. Importantly, increased *in utero* embryo accumulation can be mediated by a blend of male-released pheromones.
4. Neuropeptide control: AQR/PQR/URX pseudocoelomic cavity-opened neurons release FLP-8 and FLP-21 neuropeptides that negatively regulate exophergenesis. This type of modulation is important for the decrease in exophergenesis dependent on hermaphrodite pheromones but not for the increase in reliance on male pheromones.
5. Integrated response: Exopher production is modulated by a blend of internal processes and external cues, whether from male or hermaphrodite secretions. This balance ensures that exophergenesis is regulated in tandem with various internal and external environmental conditions.

In encapsulating our findings, the exopher regulation in *C. elegans* unfolds as a sophisticated interplay of signals, neuronal circuits, and their associated molecules. These components collectively respond and adapt to male-derived cues, and hermaphrodite-derived signals, creating a balanced system for exopher production and release (see Fig. 8 for a detailed overview).

**Figure 8.**
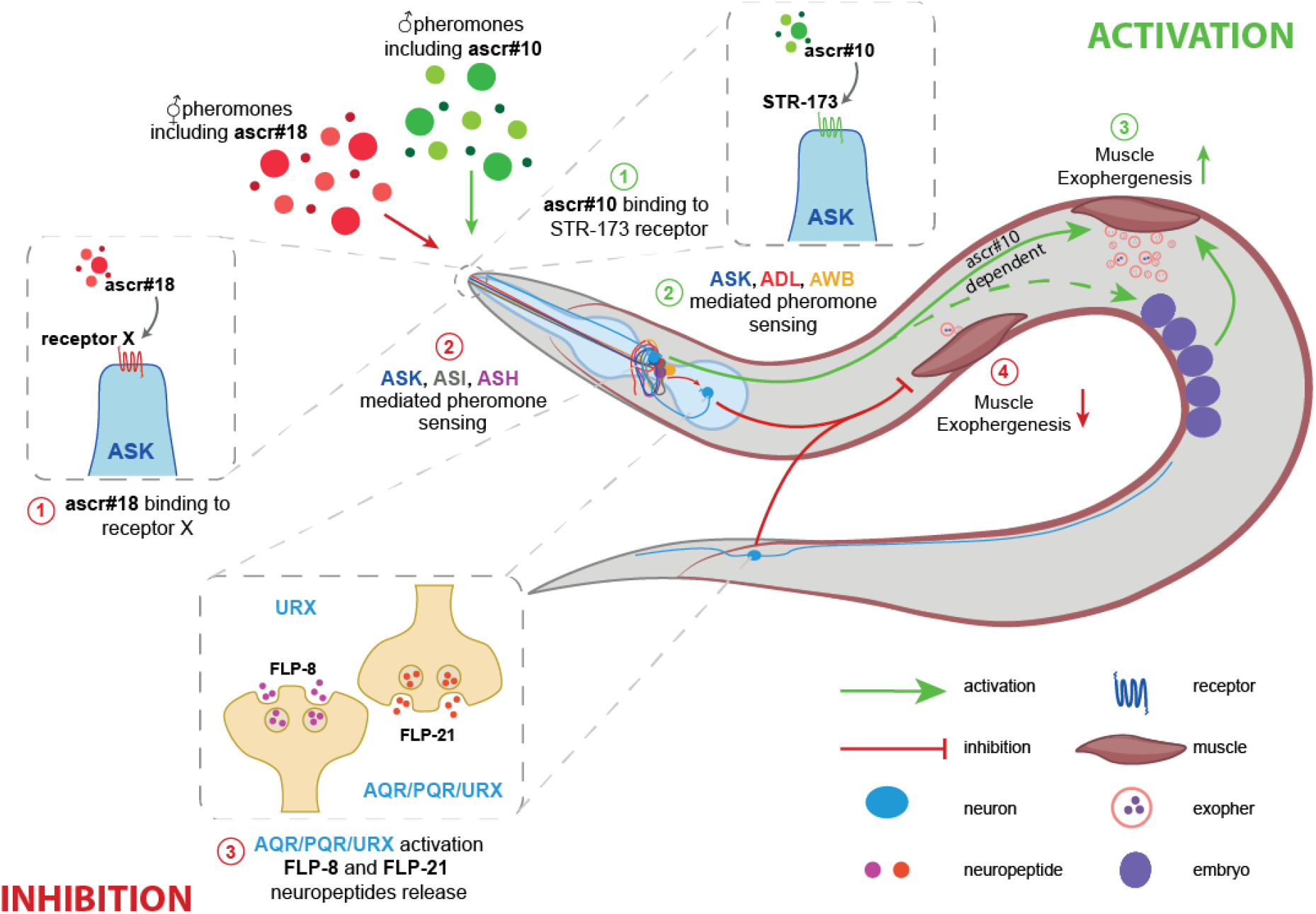
Model. Males produce ascarosides, including ascr#10, which increase exopher production through ASK, ADL, and AWB signaling. The critical player in this pathway is the ASK-expressed STR-173 G protein-coupled receptor, which enhances exopher production in response to ascr#10. In contrast, hermaphrodites release ascarosides like ascr#18, which reduce exopher production, primarily acting via ASI, ASH, and ASK (via ascr#18) sensory neurons. AQR/PQR/URX neurons in the pseudocoelomic cavity release FLP-8 and FLP-21 neuropeptides, negatively regulating exophergenesis. This modulation is crucial for decreasing exophergenesis due to hermaphrodite pheromones but not for the increase driven by male pheromones.

## Discussion

Our study shows that regulation of exophergenesis in *C. elegans* presents a sophisticated network of neuroendocrine and pheromone signaling pathways that integrate internal and external environmental cues. These results shed light on the vital roles of male- and hermaphrodite-derived ascarosides in influencing this complex network. Central to our findings is the role of male-derived ascaroside pheromones, particularly ascr#10. Detected by the STR-173 receptor in ASK neurons, this pheromone plays a pivotal role in potentiating exopher production. On the other end of the spectrum, hermaphrodites produce pheromones, including ascr#18, which act to temper exopher levels. Our investigations revealed that the ASH, ASI, and ASK olfactory neurons mediate this response. This interplay between ascaroside synthesis, chemosensation, and neuronal regulation hints at the sophisticated nature of *C. elegans*’ inter-animal communication. Our research further identified the AQR, PQR, and URX neurons as pivotal elements in the matrix of exopher regulation. The profound influence of FLP-8 and FLP-21 neuropeptides, originating from AQR, PQR, and URX neurons, further consolidated our understanding. The modulation of these peptides, particularly with their associated neurons, showcases the intricate balance of environmental and social signals with the activity of endocrine systems within the nematode.

The notable increase in exopher production among hermaphrodites either exposed to male metabolites or grown with males underscores the significant influence of male pheromones on cellular processes. This observation aligns well with prior findings suggesting male pheromones channel resource allocation towards the germline^41^. In this context, it is conceivable that exophers are a part of this mechanism, possibly playing a pivotal role in bolstering oocyte and embryo quality. This strategy may enhance reproductive success, ensuring higher quality offspring even at the potential expense of the individual’s somatic health. However, this reproductive advantage seems to come at a cost. The elevation in exophergenesis has been linked to a decreased exploratory behavior, which could be associated with swifter deterioration of muscle function with age^6^. Aprison and Ruvinsky’s observations that hermaphrodites expedite both development and somatic aging in the presence of males^39^ further solidify this notion. This underscores the importance of understanding the role and regulation of exophers, as they could serve as a nexus between environmental cues (like male pheromones), reproductive strategies, and age-related degeneration. As such, exophers probably act as biological executors and carriers of information in inter-animal communication. This is also consistent with observations that *Drosophila* males secrete EVs that are important for mating behavior^42^, and the exchange of information between male and female flies leads to increased EVs release from sperm secretory cells, which promotes fertility^43^. Drawing parallels with mammals, while the signaling molecules (like ascarosides) might differ, the broader concept of resource allocation and trade-offs between reproduction and somatic well-being might find relevance, offering avenues for future exploration.

While the proximity of males induces an increase in exophergenesis, a heightened density of hermaphrodites elicits a contrary response by diminishing it. There are instances where organisms, in response to densely populated environments, adaptively modulate their metabolic and reproductive functions to address potential resource scarcity^10,44^. Within the context of *C. elegans*, an escalation in hermaphrodite density could indicate imminent resource scarcity. Consequently, this instigates the nematodes to modify survival tactics, encompassing the reduction of exopher synthesis. Given the complexity of cues—a supportive push from embryo-maternal signals and a dampening nudge from dense hermaphrodite surroundings— the resultant exopher production could be a net outcome of these counteracting signals. Future investigations should unravel these cues’ hierarchical dynamics and identify prevailing signals under diverse conditions. For a comprehensive understanding of this facet, subsequent research should emphasize alterations in neural activity amidst different social scenarios and discerning the distinct signaling responses triggered by both male and hermaphrodite stimuli. Investigating how *C. elegans* balances the demand for supporting offspring development (via exophers) with the need to adapt to social stresses in crowded conditions can offer deeper evolutionary insights.

Our study has illuminated the multifaceted nature of the regulatory network involving ascarosides, their receptors, and olfactory neurons in the environmental adaptations of *C. elegans*. Specifically, we discovered that ascr#10, which is abundantly excreted upon sexual maturation by males, amplifies exopher production through the STR-173 receptor, whereas ascr#18, which is excreted constitutively at much lower rates by hermaphrodites, diminishes it. While the receptor for ascr#18 remains unidentified, our results indicate it must also be located within ASK neurons. This intricate interplay between chemoreceptors and pheromones suggests that exophergenesis is regulated by a complex network influenced by social and environmental cues. The impact of this network is further highlighted by our observation that the removal of an olfactory neuron expressing various chemoreceptors or alterations in the ascaroside biosynthesis pathway can either positively or negatively regulate exophergenesis.

The complexity of the entire regulatory network is escalated by the fact that the lack of activity of one receptor, achieved either by transcriptional downregulation or post-translational modification, can drastically alter the biological potential of its ligands. This was demonstrated in our analysis, where knockdown of *str-173* reversed the pro-exopher potential of ascr#10, implying that even subtle alterations in receptor activity can have profound consequences on the physiological responses mediated by the associated ligands. Identifying the anti-exopher factors will be pivotal in understanding the full spectrum of regulatory effects mediated by ascarosides and their receptors. Additionally, our study exemplifies the multilayered interplay between pheromones and cellular and organismal responses. The notable accumulation of embryos in the hermaphrodite’s uterus upon exposure to the male secretome, contrasting with the lack of egg retention induced by ascr#10, underlines the intricate connection between external pheromonal cues and internal physiological responses, suggesting a strategic evolutionary adaptation for resource allocation in response to reproductive efforts. These findings also highlight that ascr#10 is likely one of several male-excreted small molecule signals that affect exopher formation and egg laying in hermaphrodites. Moreover, the interaction of ascr#10- and ascr#18-mediated signaling with each other and additional male-derived signals can reveal additional pathways and responses. In conclusion, our findings underscore the need for extensive research into the complex interplay between metabolites (pheromones/ascarosides), receptors, and intracellular processes and their modulation by social and environmental cues. Understanding this network will deepen our knowledge of the mechanisms governing exophergenesis and provide valuable insights into other cellular processes and environmental adaptations mediated by similar mechanisms in *C. elegans* and potentially other organisms.

Our research demonstrates the pivotal role of the STR-173 receptor, specifically in ASK neurons, in orchestrating pheromone-driven exophergenesis. This discovery contributes to the limited knowledge surrounding 7TM GPCRs as ascaroside/pheromone receptors, considering the vast number of 7TM GPCRs expressed^45^ and metabolites secreted by *C. elegans*^46^. The association between STR-173 receptors and male ascr#10 suggests a receptor-ligand dynamic, potentially triggering a series of intracellular processes that modulate exopher production. A comprehensive exploration of this cascade, spanning receptor engagement to downstream effects, remains elusive. In the intricate environment of *C. elegans*, our investigation has provided illuminating insights into the multifaceted roles played by specific sensory neurons AQR, PQR, and URX and their associated neuropeptides, FLP-8 and FLP-21, in modulating muscle exopher production. Building on prior studies, our identification of FLP-8 and FLP-21 as modulators of extracellular vesicle release from body wall muscles brings further depth to our understanding of neuropeptide functionality in *C. elegans*.

Neuropeptides, exemplified by FLP-18, demonstrate intricate roles in regulating muscle function and behavior in *C. elegans*. Specifically, the co-release of FLP-18 with tyramine from the RIM neurons and its targeting of body wall muscles via the NPR-5 receptor to modulate muscle excitability underscores the multifaceted interactions of neuropeptides and neurotransmitters in these processes^47^. Given this, it is plausible to assume that FLP-8 and FLP-21 may function along similar pathways. Their involvement might influence muscle excitability in a manner that subsequently impacts the release dynamics of extracellular vesicles. Exploring the precise receptors and pathways through which FLP-8 and FLP-21 act will be instrumental in further elucidating their specific roles in exophers release and muscle functionality.

In *C. elegans*, specific neuropeptides like FLP-8 and FLP-21, along with distinct neuronal classes such as AQR, PQR, and URX, appear to play a pivotal role in the modulation of exopher production. Given that exophers encapsulate damaged or aggregated proteins, understanding the regulatory mechanisms in this nematode model might offer a novel perspective on protein aggregation and its clearance. While these specific neuropeptides and neuronal populations are unique to *C. elegans*, the underlying principles might hold broader biological significance. For example, if analogous mechanisms exist in more complex organisms, they could potentially impact our understanding of diseases like Alzheimer’s or Parkinson’s, where protein aggregation is a prominent concern. Thus, while direct translational insights may be limited, *C. elegans* offers a window into fundamental processes that could shed light on the sophisticated dance between neurons, protein aggregation, and potential clearance mechanisms in neurodegenerative conditions.

In conclusion, our research is an initial foray into unraveling the complex regulatory mechanisms of exopher production in *C. elegans*, suggesting the possibility of universally conserved physiological systems. The findings hint at a future filled with deeper exploration into these dynamics, possibly redefining our understanding of intercellular communication across species. Similar mechanisms modulating cell-to-cell communications via exophers might exist in humans, as evidenced by studies on other extracellular vesicle classes^43^. Our study also underscores the significance of the worm’s olfactory system in steering exopher generation within muscles. It is worth noting that human olfactory impairments are linked with a heightened risk for cardiovascular disease (CVD)^44,45^. Such a correlation might be attributed to disruptions in exopher-driven homeostatic processes within the heart. Still, it remains imperative to ascertain if a parallel olfactory-driven exopher regulatory mechanism exists in mammals.

## Figures and Figure legends

**Supplementary Figure 1.**
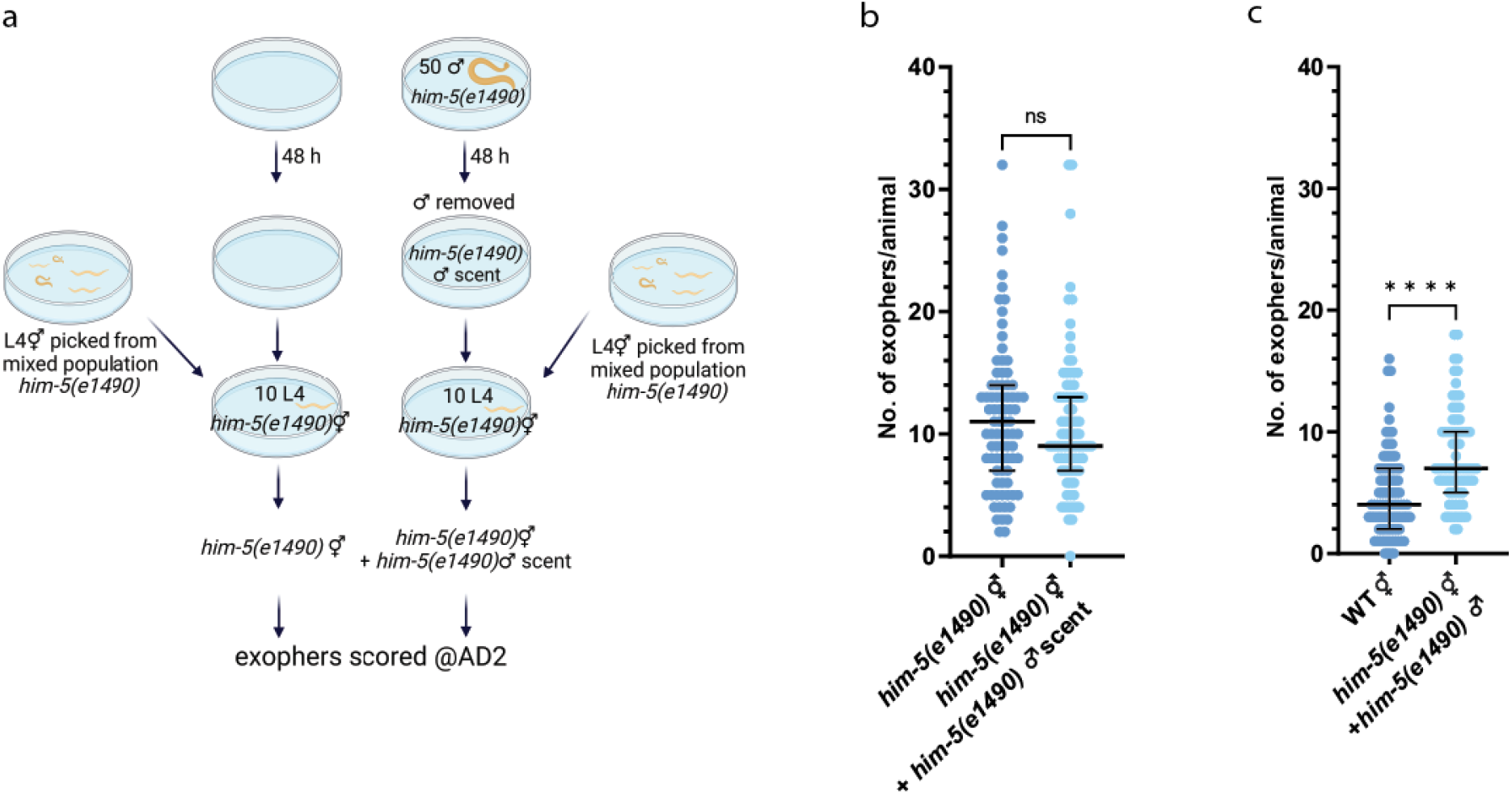
Exposing hermaphrodites to males or their scent throughout the entire duration of the experiment do not further elevate exopher production. **a** Schematic representation of the experimental setup for Supplementary Fig. 1b. **b** Exposing hermaphrodites to a male’s secretome after the L4 stage does not further increase exopher production. n = 83; N = 3. **c** Co-culturing him-5 hermaphrodites with males increases exophergenesis in hermaphrodites. n = 72 and 90; N = 3. Data information: Data are presented as median with interquartile range; n represents the number of worms; N represents a number of experimental repeats that were combined into a single value; ns - not significant (P > 0.05), **** P < 0.0001; Mann-Whitney test.

**Supplementary Figure 2.**
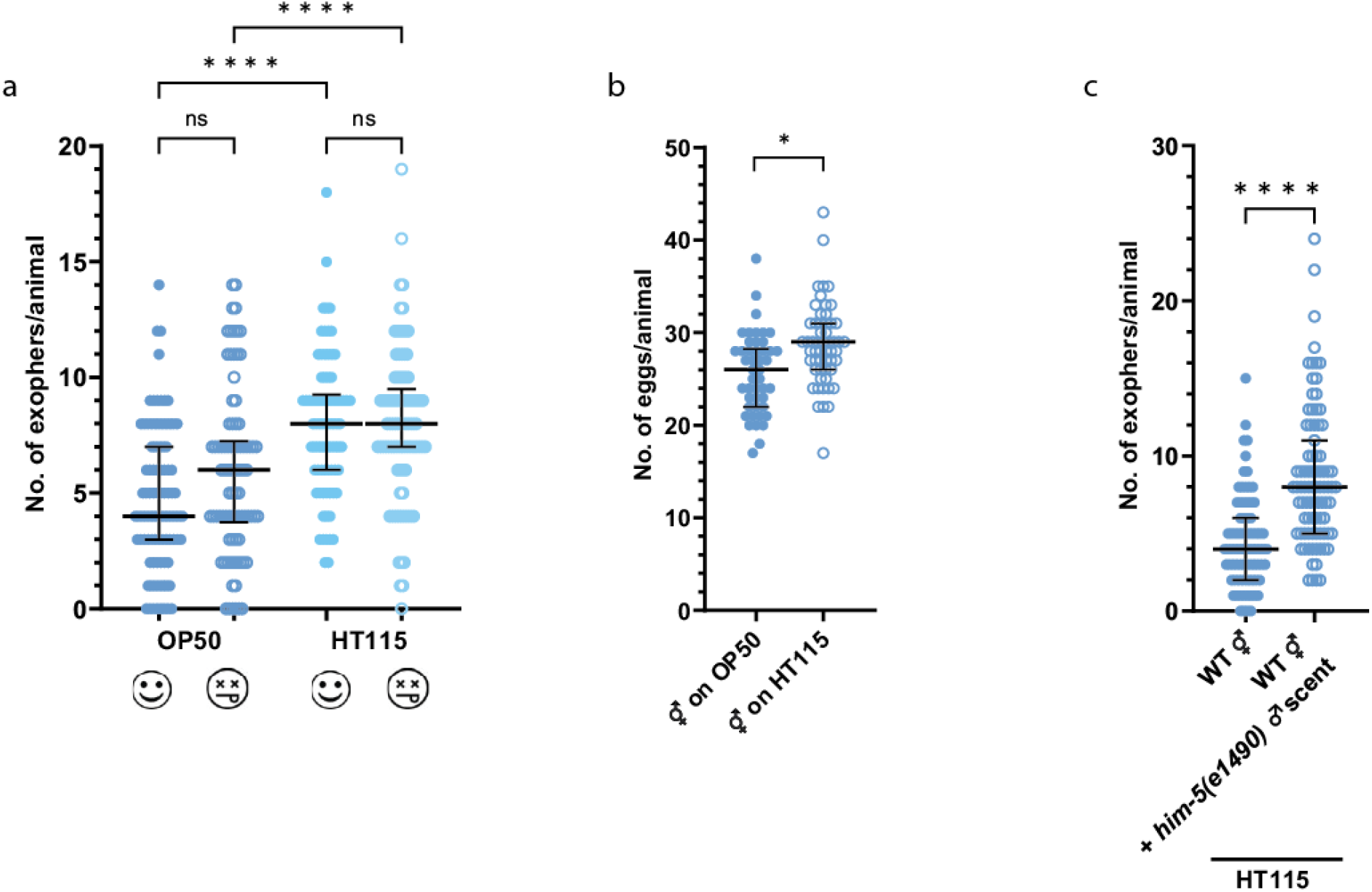
Influence of bacteria diet on muscle exophers formation. **a** Worms fed with metabolically active and inactive bacteria produce a similar number of exophers. Worms grown on *E. coli* HT115 strain produce more exophers than worms grown on *E. coli* OP50 strain. n = 78 - 93; N = 3. **b** Worms grown on *E. coli* HT115 strain contain more eggs *in utero* than worms grown on *E. coli* OP50 strain. n = 46 and 55; N = 3. **c** Growing hermaphrodites on male-conditioned plates increases exophergenesis levels regardless of *E. coli* strain used as a food source. n = 79 and 87; N = 3. Data information: Data are presented as median with interquartile range; n represents the number of worms; N represents a number of experimental repeats that were combined into a single value; ns - not significant (P > 0.05), * P < 0.05, **** P < 0.0001; (a) Kruskal-Wallis test with Dunn’s multiple comparisons test, (b, c) Mann-Whitney test.

**Supplementary Figure 3.**
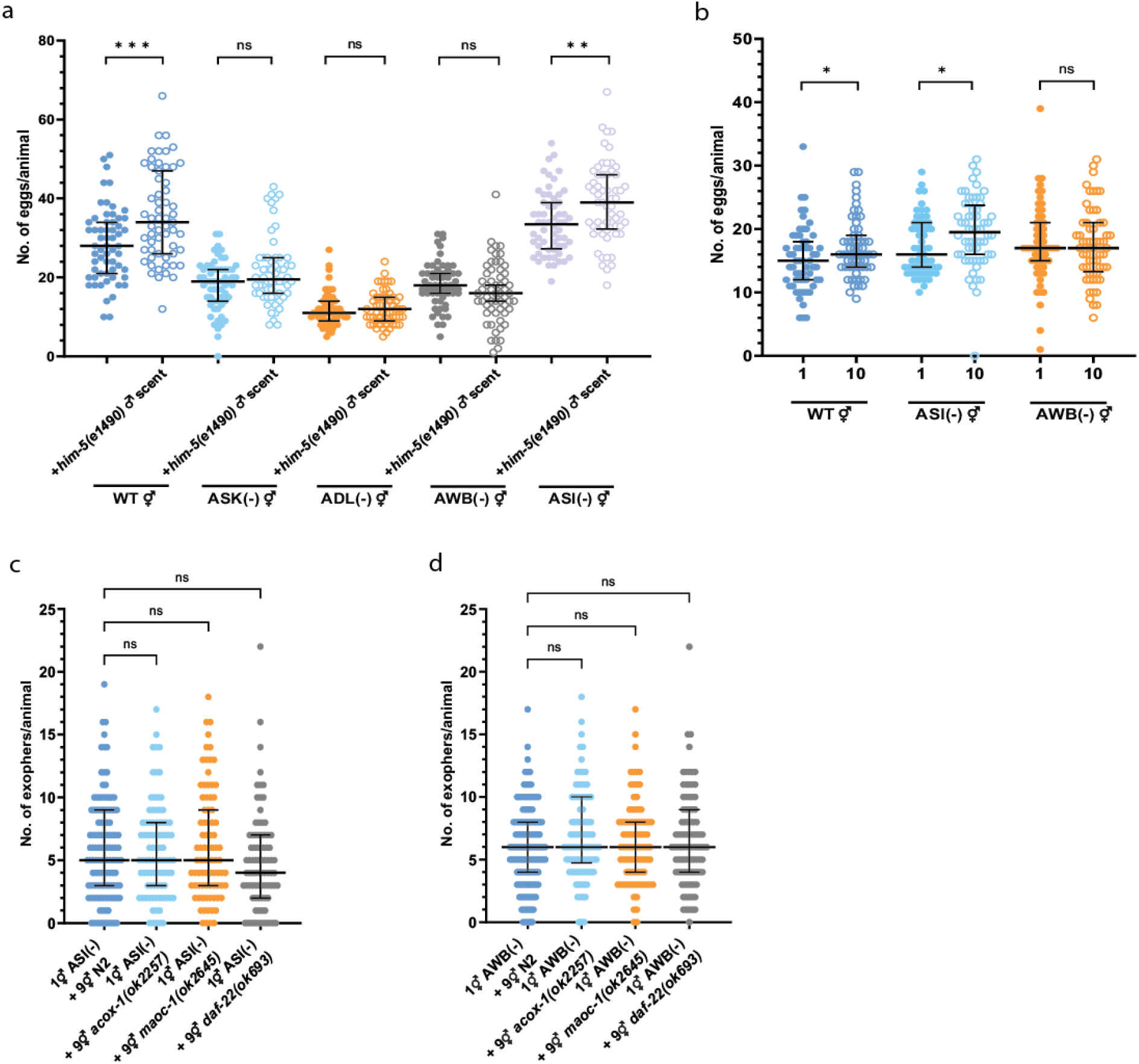
Influence of pheromones released by ascaroside side-chain biosynthesis mutants on hermaphrodites with genetically ablated sensory neurons. **a** Removal of ASK, AWB, or ADL neurons prevented the increase in the number of embryos in utero after exposing hermaphrodites to male pheromones. n = 55 - 60; N = 3. **b** *In utero* embryo number for wild-type worms and ASI and AWB ablation mutants grown as solitary animals or in ten-hermaphrodites population. n = 55 - 60; N = 3. **c - d** Worms with ablated ASI and AWB neurons grown together with ascaroside side-chain biosynthesis mutants do not exhibit alterations in produced exopher numbers. n = 83 - 120; N = 3. Data information: Data are presented as median with interquartile range; n represents the number of worms; N represents a number of experimental repeats that were combined into a single value; ns - not significant (P > 0.05), * P < 0.05, ** P < 0.01,*** P < 0.001; (a, b) Mann-Whitney test, (c, d) Kruskal-Wallis test with Dunn’s multiple comparisons test.

**Supplementary Figure 4.**
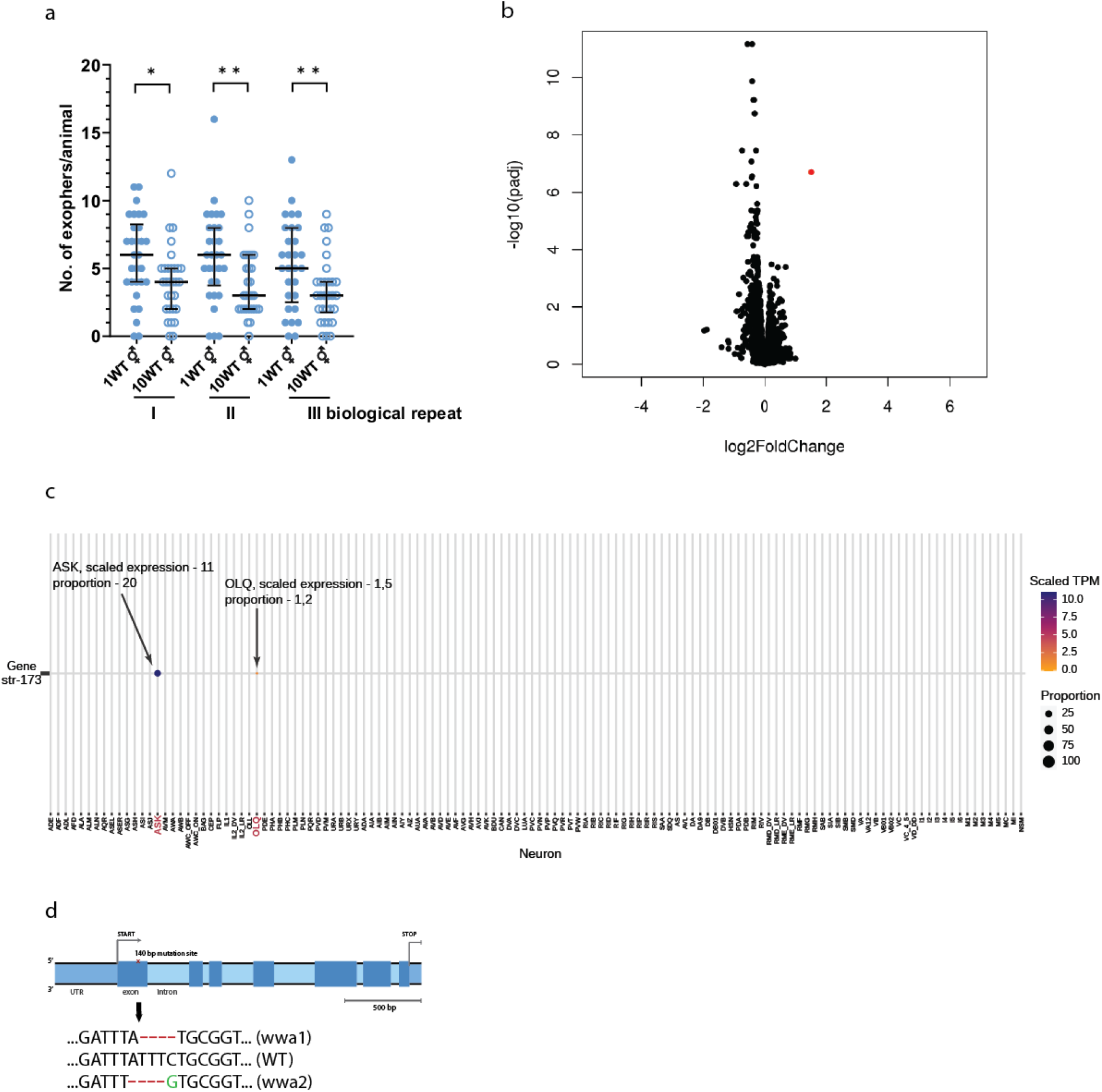
Worms grown in populations with different densities did not differ significantly on the transcriptional level. **a** Number of exophers produced by worms that were used for RNAseq analysis. n = 29 - 30; N = 3. **b** Worms grown as a single animal or in a ten-hermaphrodites population did not differ significantly on the transcriptional level. A transcript with significant change is marked as a red dot. **c** Single-cell RNA-seq data from CeNGENApp^35^ show *str-173* strong expression in ASK neurons and weak expression in OLQ neurons. The circle diameter represents the proportion of neurons in each cluster that express the *str-173* gene. **d** Localization of *str-173* mutations in the gene. Data information: Data are presented as median with interquartile range; n represents the number of worms; N represents a number of experimental repeats that were combined into a single value; * P < 0.05, ** P < 0.01; (**a**) Mann-Whitney test.

**Supplementary Figure 5.**
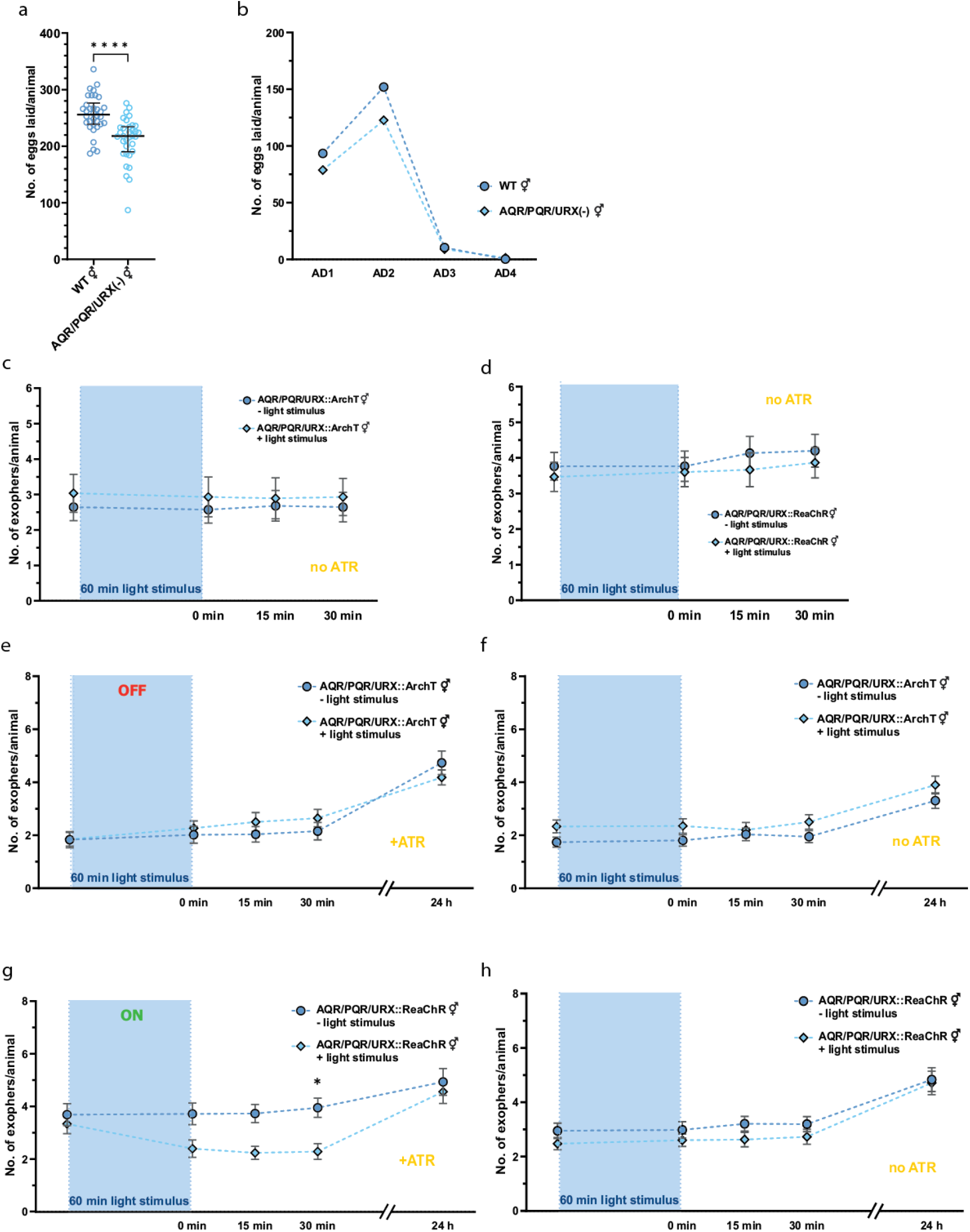
ReaChr-based activation but not ArchT-based inactivation at the adult day 1 (AD1) stage influence exopher production in hermaphrodites. **a - b** Worms with the genetic ablation of pseudocoelom-exposed AQR, PQR, and URX neurons have lower brood size than the wild type. n = 34; N = 3. **c - d** Control experiment without all-trans retinal (ATR) for ArchT-mediated inactivation and ReaChR-mediated activation of AQR, PQR, and URX neurons. n = 30; N = 3. **e – f** ArchT-mediated inactivation of AQR, PQR, and URX neurons at the adult day 1 (AD1) stage do not modulate exopher production. n = 59 - 72; N = 5 - 6. **g – h** ReaChR-mediated activation of AQR, PQR, and URX neurons at the adult day 1 (AD1) stage significantly decreases exopher production. n = 59 - 62; N = 5 - 6. Data information: +ATR means “with all-trans-retinal”; no ATR means “without all-trans-retinal”. Data are presented as median with interquartile range (a), mean (b), or mean with SEM (c – h); n represents the number of worms; N represents a number of experimental repeats that were combined into a single value; ns - not significant * P < 0.05, **** P < 0.0001; Mann-Whitney test.

**Supplementary Figure 6.**
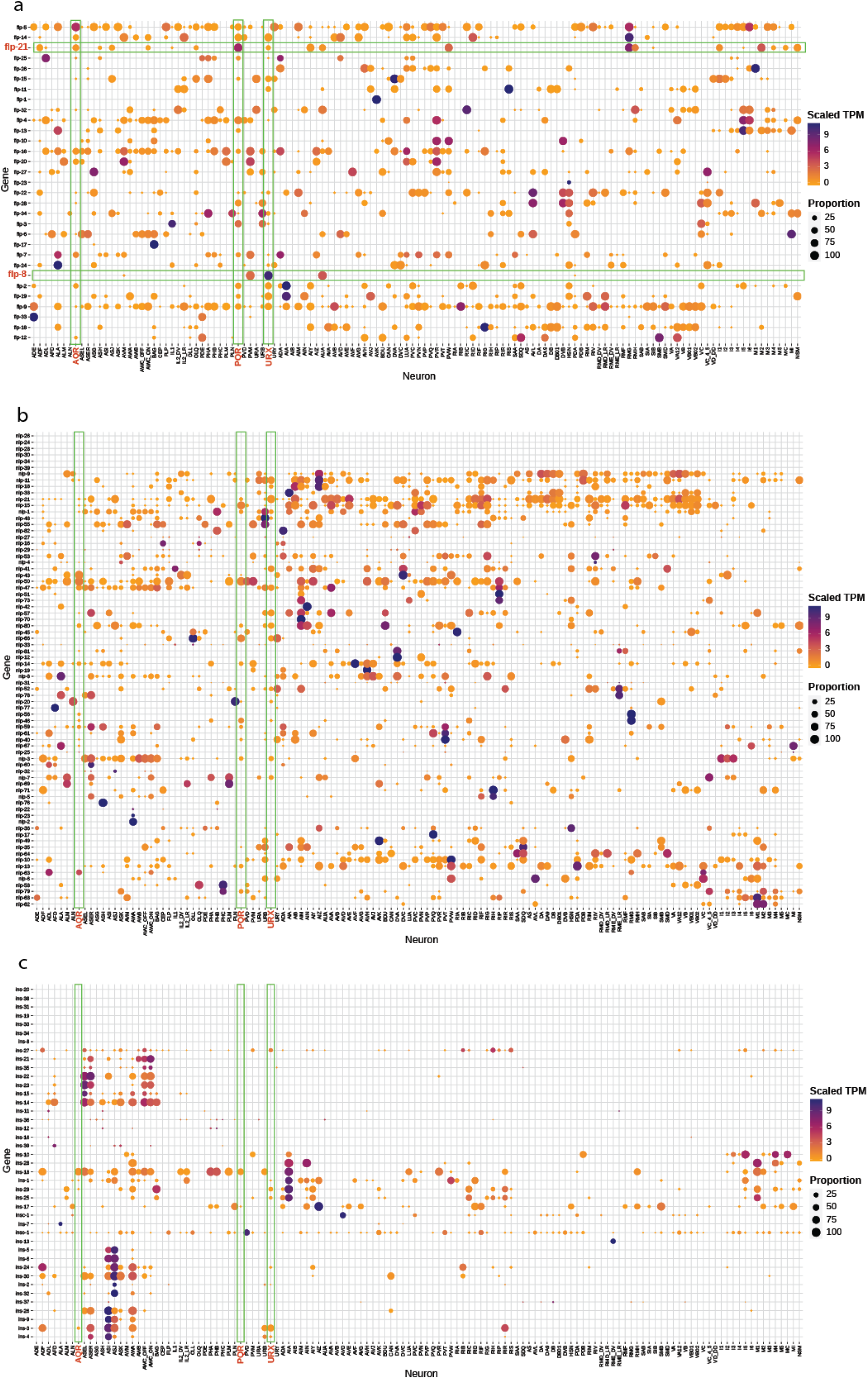
FLP-8 and FLP-21 are expressed in URX and AQR, PQR, URX, respectively. **a - c** *flp-8* and *flp-21* are strongly and relatively specifically expressed in AQR, PQR and/or URX neurons. Data taken from^35^.

**Supplementary Figure 7.**
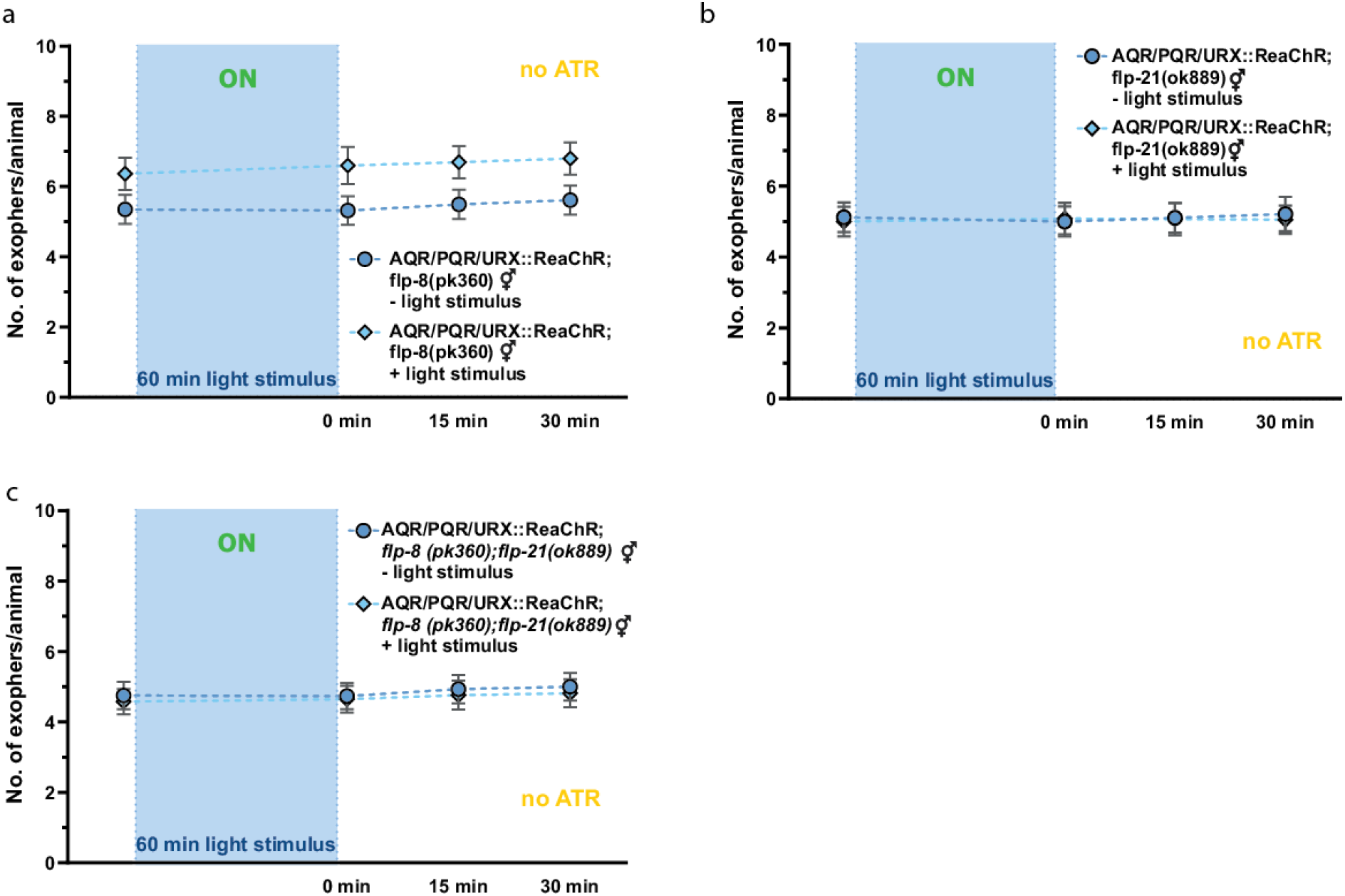
Control experiments without all-trans retinal (ATR) ReaChR-mediated activation of AQR, PQR, and URX neurons that do not produce FLP-8 and/or FLP-21 neuropeptides. **a** Control experiment without all-trans retinal (ATR) ReaChR-mediated activation of AQR, PQR, and URX neurons in *flp-8(pk360)* mutant background. n = 60; N = 5. **b** Control experiment without all-trans retinal (ATR) ReaChR-mediated activation of AQR, PQR, and URX neurons in *flp-21(ok889)* mutant background. n = 57 and 58; N = 5. **c** Control experiment without all-trans retinal (ATR) ReaChR-mediated activation of AQR, PQR, and URX neurons in *flp-8(pk360), flp-21(ok889)* double mutant background. n = 59 and 60; N = 5. Data information: +ATR means “with all-trans-retinal”; no ATR means “without all-trans-retinal”. Data are presented as mean with SEM; n represents the number of worms; N represents a number of experimental repeats that were combined into a single value.

---

**Supplementary Table 1.**
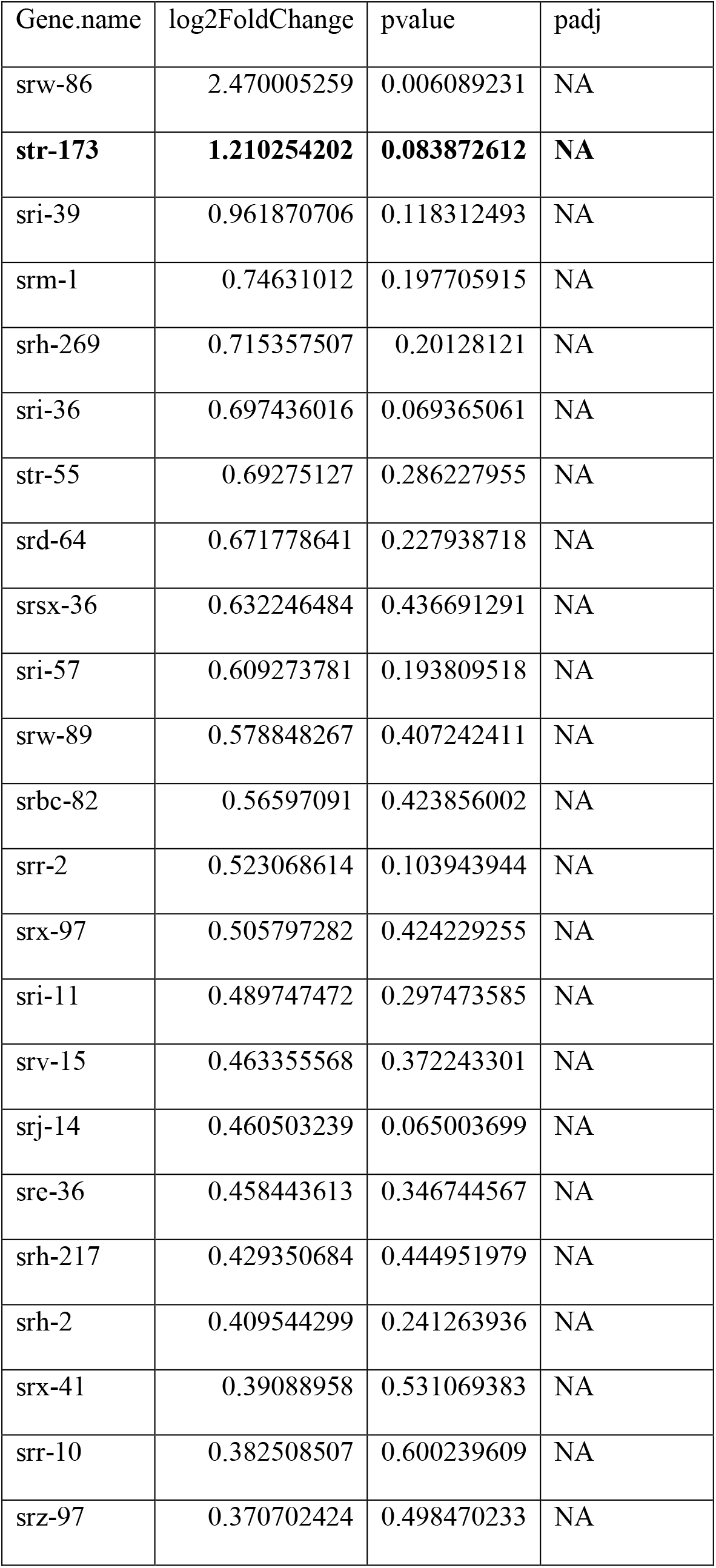
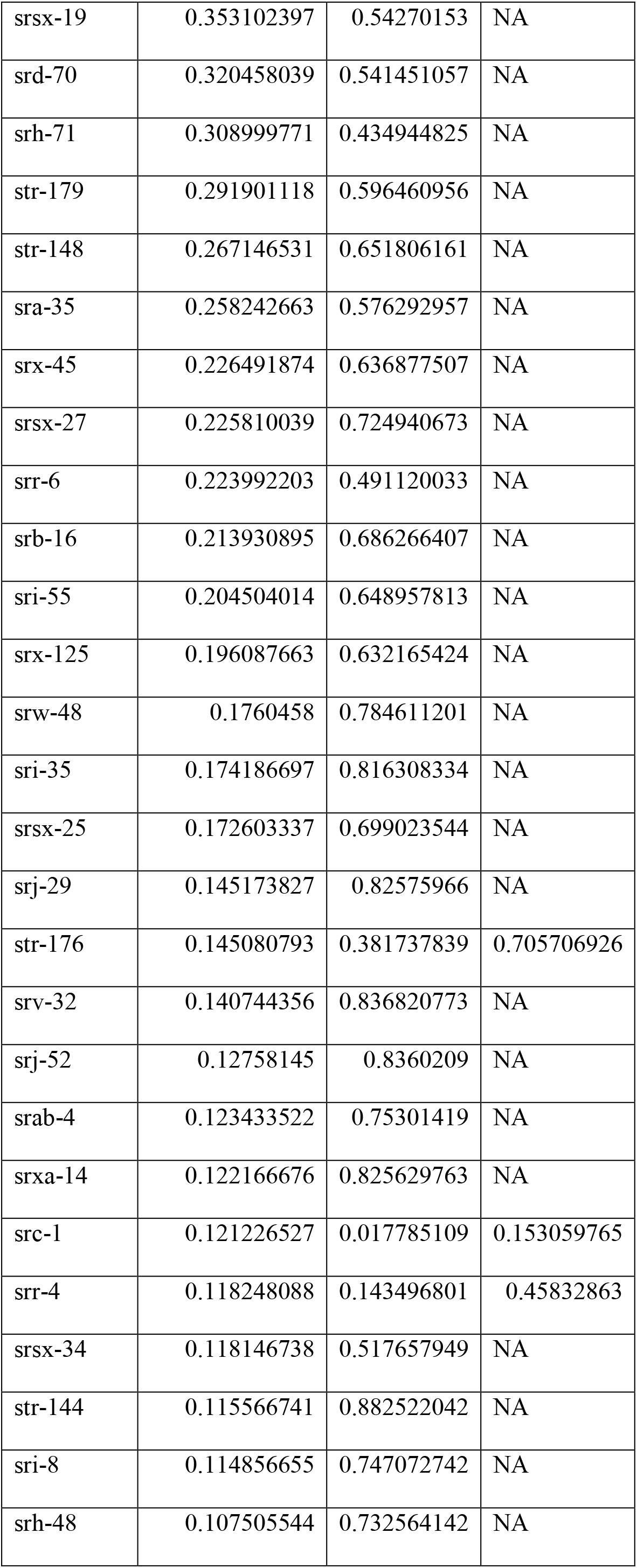
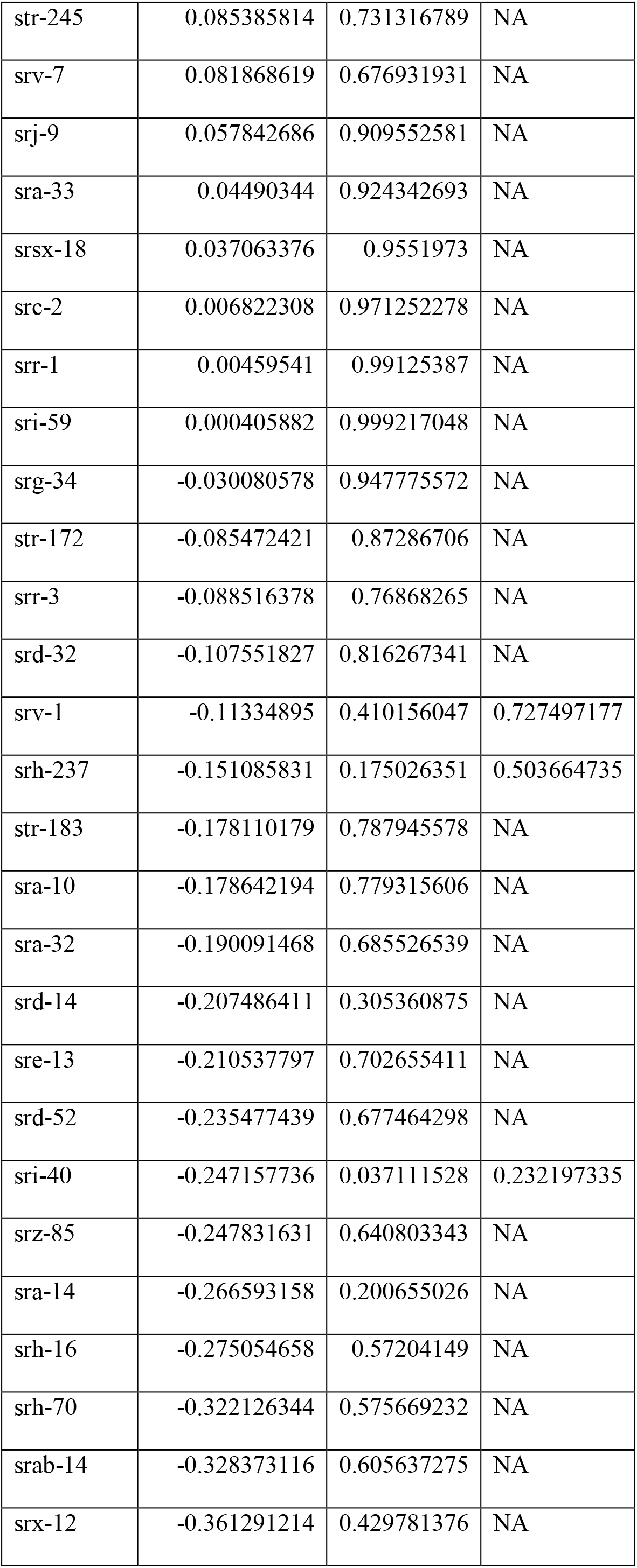
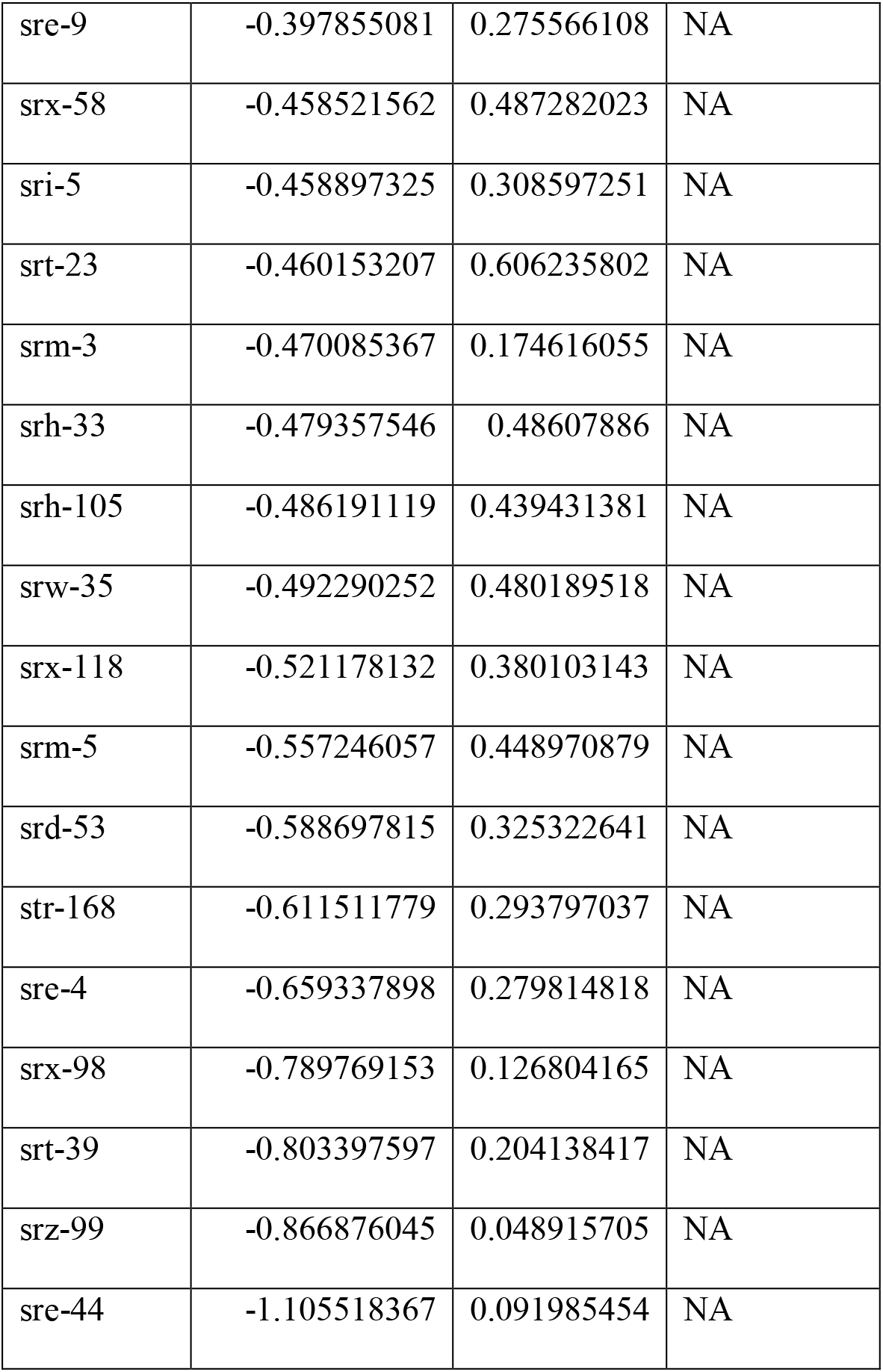

**Supplementary Table 2.** List of all exopher-related phenotypes presented in the manuscript.

| Figure no. | Condition | Observed phenotype |
| --- | --- | --- |
| 1c | Exopher level in hermaphrodites in the presence of *him-5(e1490)* males | Increased exopher number in *him-5(e1490)* mutant hermaphrodites |
| 1d | *In utero* embryo retention in the presence of *him-5(e1490)* males | Increased embryo retention in *him-5(e1490)* mutant hermaphrodites |
| 1f | Effect on exopher production in hermaphrodites grown on WT and *him-5* males-conditioned plates | Increased exopher number after exposure to *him-5* and WT males secretome |
| 1g | Effect on *in utero* embryo retention in hermaphrodites grown on WT and *him-5* males-conditioned plates | Increased *in utero* embryo retention after exposure to *him-5* and WT males secretome |
| 1h | Effect of male pheromones presence on exopher production in different developmental time frames | Increased exopher number (strongest effect on exposure during L4 to AD1 stages) |
| 1i | Effect of male pheromones presence on *in utero* embryo retention in different developmental time frames effect | Increased *in utero* embryo retention (apart from exposure during L4 to AD1 stages) |
| 2b | Population size effect on exopher production (single or 10 hermaphrodites) | Single worms produce more exophers than worms cultured in 10 hermaphrodite population |
| 2c | Population size effect on *in utero* embryo retention | No effect |
| 2d | Population size effect on exopher production (single, 5 or 10 hermaphrodites) | Exopher number decreased in 5 and 10 hermaphrodites populations |
| 2e | Population size effect on exopher production (single, 10 or 100 hermaphrodites) | Exopher number decreased in 10 and 100 hermaphrodite populations |
| 2f | Hermaphrodites-conditioned plates effect on exopher production | Hermaphrodite pheromones similarly decrease exopher number as hermaphrodite presence |
| 2g | Larvae presence effect on exopher production | Larvae presence decreases exopher production |
| 3c | Influence of ascaroside side-chain biosynthesis mutants presence on exophergenesis level in wild-type worms | Number of exophers elevated in the presence of *acox-1* mutants; decreased exopher level in the presence of *maoc-1* mutants; *daf-22* mutants presence does not affect exophergenesis |
| 3d | Influence of ascaroside side-chain biosynthesis mutants presence on *in utero* embryo retention | No influence |
| 3e | Effect of ascr#10 exposure on exophergenesis | Dose-dependent effect– increase in exopher production after 1ng ascr#10 exposure; exposure to 1 pg ascr#10 has no effect |
| 3f | Effect of ascr#10 on *in utero* embryo retention | No effect |
| 3g | Effect of ascr#18 exposure on exophergenesis | Decrease in exopher number after exposure to 1pg or 1ng of ascr#18 |
| 3h | Effect of ascr#18 on *in utero* embryo retention | Decreased in utero retention after exposure to 1ng of ascr#18 |
| 4b | Influence of impaired ciliated sensory neurons on exophergenesis in *che-13* mutant | *che-13* mutant produces fewer exophers than wild-type worms |
| 4c | Genetic ablation of chemosensory neurons effect on exopher production | ASK(-), AWC(-), ADL(-), or ASH(-) mutants produce fewer exophers |
| 4d | Genetic ablation of chemosensory neurons effect on *in utero* embryo retention | ASK(-), AWC(-), ADL(-), ASH(-) and CEM(+) have decreased *in utero* embryo retention |
| 4e | Influence of AWC neuron absence on exophergenesis in different temperatures | AWC neurons regulate the temperature-dependent increase in muscle exopher production |
| 4f | Male-specific pheromones effect on exophergenesis in mutants with ablated chemosensory neurons | ASK(-), ADL(-), AWB(-) – no effect<br>WT, AWC(-), ASH(-), ASI(-), CEM(+) – increased exophergenesis after male pheromones exposure |
| 4g | Population size effect on exopher production in mutants with ablated chemosensory neurons | ASH(-), ASI(-), CEM(-), ASK(-) – no effect<br>AWC(-), ADL(-), AWB(-) – exopher number decreased in worms grown in the population of 10 hermaphrodites |
| 5a | *str-173* expression visualization in L1, L4 and AD2 worms | *str-173* is expressed in ASK neurons, vulva, and the tail in L4 and adult worms;<br>L1 larvae express *str-173* in pharynx |
| 5b | *str-173* expression visualization in L4 worms | *str-173* is expressed in ASK neurons |
| 5c | Exopher level in *str-173* mutants | Exopher production is not altered in *str-173* mutants |
| 5d | Population size effect on exopher production in *str-173* mutants | *str-173(wwa1)* mutation does not affect exopher production in worms grown in separation or in 10 hermaphrodites population;<br>*str-173(wwa2)* – higher exopher level in worms grown in separation |
| 5e | Male-specific pheromones effect on exopher production in *str-173* mutants | No effect |
| 5f | Effect of ascr#10 exposure on exophergenesis in *str-173* mutants | Decrease in exopher production |
| 5g | Effect of ascr#10 on eggs in utero retention in *str-173* mutants | Decreased *in utero* embryo retention |
| 5h | Effect of ascr#18 exposure on exophergenesis in *str-173* mutants | Decrease in exopher production |
| 5i | Effect of *ascr#18* on eggs in utero retention in *str-173* mutants | Decreased *in utero* embryo retention |
| 5j | Effect of *ascr#10* and *ascr#18* exposure on exophtogenesis in ASK(-) mutant | No effect |
| 6a | Ablation of AQR/PQR/URX neurons effect on exopher level | Increased exopher number in AQR/PQR/URX(-) mutant |
| 6b | Ablation of AQR/PQR/URX neurons effect *in utero* embryo retention | Decreased *in utero* embryo retention in AQR/PQR/URX(-) mutant |
| 6c | ArchT-mediated AQR/PQR/URX inactivation effect on exopher production (AD2) | Increased exopher number after AQR/PQR/URX inactivation |
| 6d | ReaChR-mediated AQR/PQR/URX activation effect on exopher production (AD2) | Decreased exopher number after AQR/PQR/URX activation |
| 6e | Ablation of ASK/AQR/PQR/URX neurons effect on exopher level | Low exophtogenesis levels in animals with genetic ablation of ASK neurons and high levels of exophtogenesis in animals with genetic ablation of AQR, PQR, and URX neurons are equalized in animals with all four neurons removed |
| 6f | Population size effect on exopher production in AQR/PQR/URX(-) mutant | No effect |
| 6g | Male-specific pheromones effect on exopher production in AQR/PQR/URX(-) mutant | Increase in exopher production after exposure to male secretome |
| 6h | FUdR effect on exophtogenesis in AQR/PQR/URX(-) mutant | Decreased exopher production in animals treated with FUdR is not rescued by AQR/PQR/URX removal |
| 6i | *rme-2* knockdown effect on exophtogenesis in AQR/PQR/URX(-) mutant | AQR/PQR/URX removal only partially rescues the inhibition of exophtogenesis caused by *rme-2* knockdown |
| 7a | Exopher level in *flp-8* and *flp-21* mutants | Increased exopher production in single *flp-8* and *flp-21* mutants, and in *flp-8; flp-21* double mutant |
| 7b | Number of *in utero* embryos in *flp-8* and *flp-21* mutants | No change |
| 7c | ReaChR-mediated AQR/PQR/URX; *flp-8(tm741)* activation effect on exopher production (AD2) | Increased exopher production |
| 7d | ReaChR-mediated AQR/PQR/URX; *flp-21(ok889)* activation effect on exopher production (AD2) | No effect |
| 7e | ReaChR-mediated AQR/PQR/URX; *flp-8(tm741); flp-21(ok889)* activation effect on exopher production (AD2) | No effect |
| Sup. Fig. 1b | *him-5* males secretions effect on *him-5* hermaphrodites' exopher production | No effect |
| Sup. Fig. 1c | *him-5* males presence effect on *him-5* hermaphrodites eggs in utero retention | Increased number of *in utero* embryo retention when *him-5* males are present |
| Sup. Fig. 2a | Food source (metabolically active and inactive bacteria) effect on exopher production | Worms fed with metabolically active and inactive bacteria produce a similar number of exophers |
| Sup. Fig. 2b | Food source (*E. coli* strain OP50 and HT115) effect on *in utero* embryo retention | More embryos were retained in the uterus when grown on HT115 |
| Sup. Fig. 2c | Males-conditioned plates seeded with HT115 bacteria strain effect on exopher production | Increase in exopher production |
| Sup. Fig. 3a | Male-specific pheromones effect on egg retention in mutants with ablated chemosensory neurons | ASI(-) – increased number of *in utero* embryos after exposure to male pheromones<br>ASK(-), ADL(-), AWB(-) – no change |
| Sup. Fig. 3b | Population size effect on *in utero* embryos retention in mutants with ablated chemosensory neurons | ASI(-) -more embryos retained in worms grown in the population of 10 hermaphrodites<br>AWB(-) – no change |
| Sup. Fig. 3c | Influence of ascaroside side-chain biosynthesis mutants presence on exophogenesis level in ASI(-) mutants | No influence |
| Sup. Fig. 3d | Influence of ascaroside side-chain biosynthesis mutants presence on exophogenesis level in AWB(-) mutants | No influence |
| Sup.<br>Fig. 4a | Population size effect on the number of exophers produced by worms that were used for RNAseq analysis | Worms cultured in a population of 10 hermaphrodites – fewer exophers produced in each biological repeat |
| Sup.<br>Fig. 5a | Number of eggs laid by AQR/PQR/URX(-) mutant | Decreased number of laid eggs by AQR/PQR/URX(-) mutant |
| Sup.<br>Fig. 5b | Number of eggs laid by AQR/PQR/URX(-) mutant on consecutive days of adulthood | No significant change |
| Sup.<br>Fig. 5c | ArchT-mediated AQR/PQR/URX inactivation effect on exopher production (no ATR control) | No change |
| Sup.<br>Fig. 5d | ReaChR-mediated AQR/PQR/URX activation effect on exopher production (no ATR control) | No change |
| Sup.<br>Fig. 5e | ArchT-mediated AQR/PQR/URX inactivation effect on exopher production (AD1) | No change |
| Sup.<br>Fig. 5f | ArchT-mediated AQR/PQR/URX inactivation effect on exopher production (no ATR control; AD1) | No change |
| Sup.<br>Fig. 5g | ReaChR-mediated AQR/PQR/URX activation effect on exopher production (AD1) | Decrease in exopher production after 30 min light stimulation |
| Sup.<br>Fig. 5h | ReaChR-mediated AQR/PQR/URX activation effect on exopher production (no ATR control, AD1) | No change |
| Sup.<br>Fig. 7a | ReaChR-mediated AQR/PQR/URX; *flp-8(tm741)* activation effect on exopher production (no ATR control) | No change |
| Sup.<br>Fig. 7b | ReaChR-mediated AQR/PQR/URX; *flp-21(ok889)* activation effect on exopher production (no ATR control) | No change |
| Sup.<br>Fig. 7c | ReaChR-mediated AQR/PQR/URX; *flp-8(tm741); flp-21(ok889)* activation effect on exopher production (no ATR control) | No change |

## Materials and Methods

### Reagents and Tools table

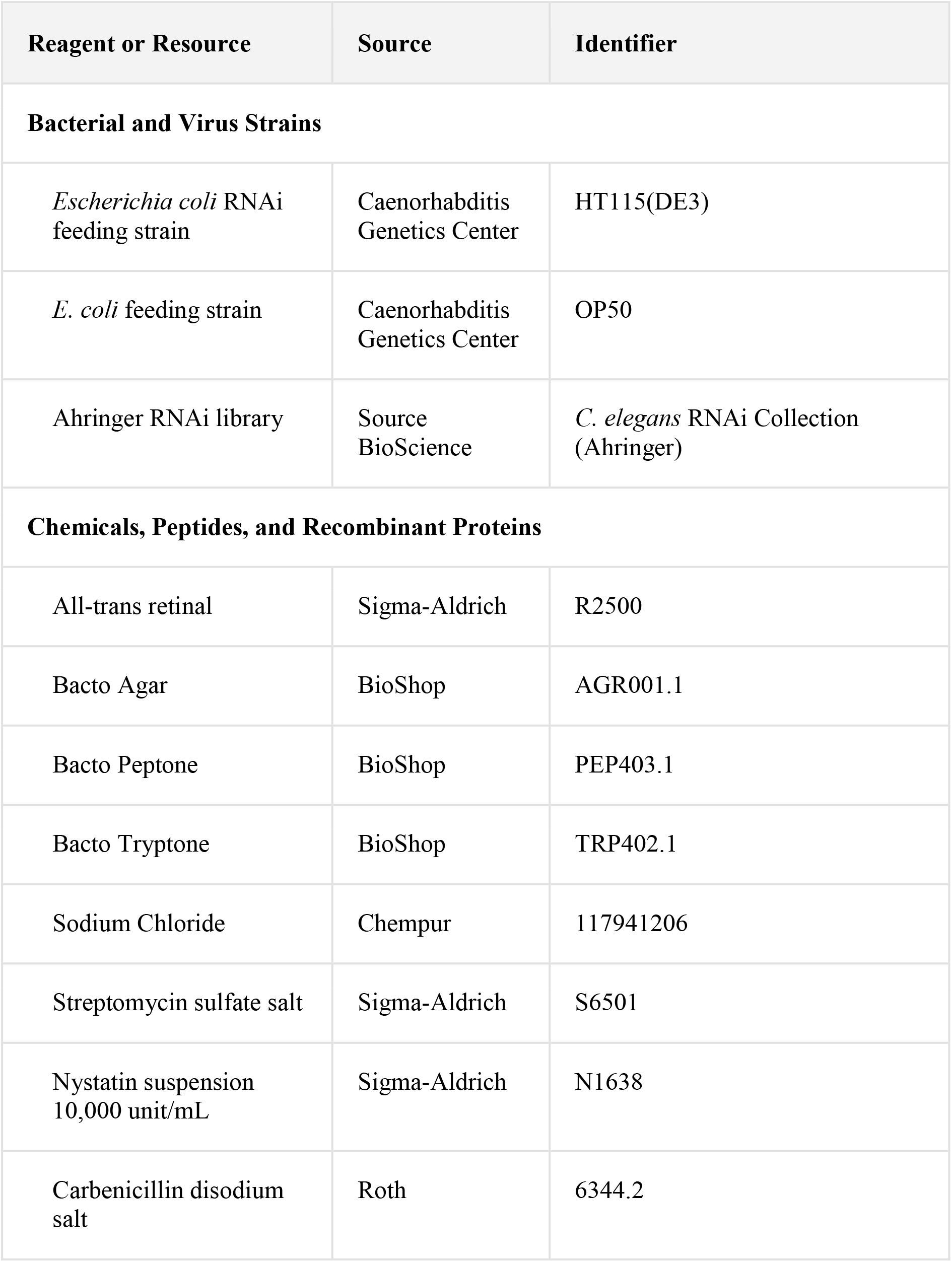
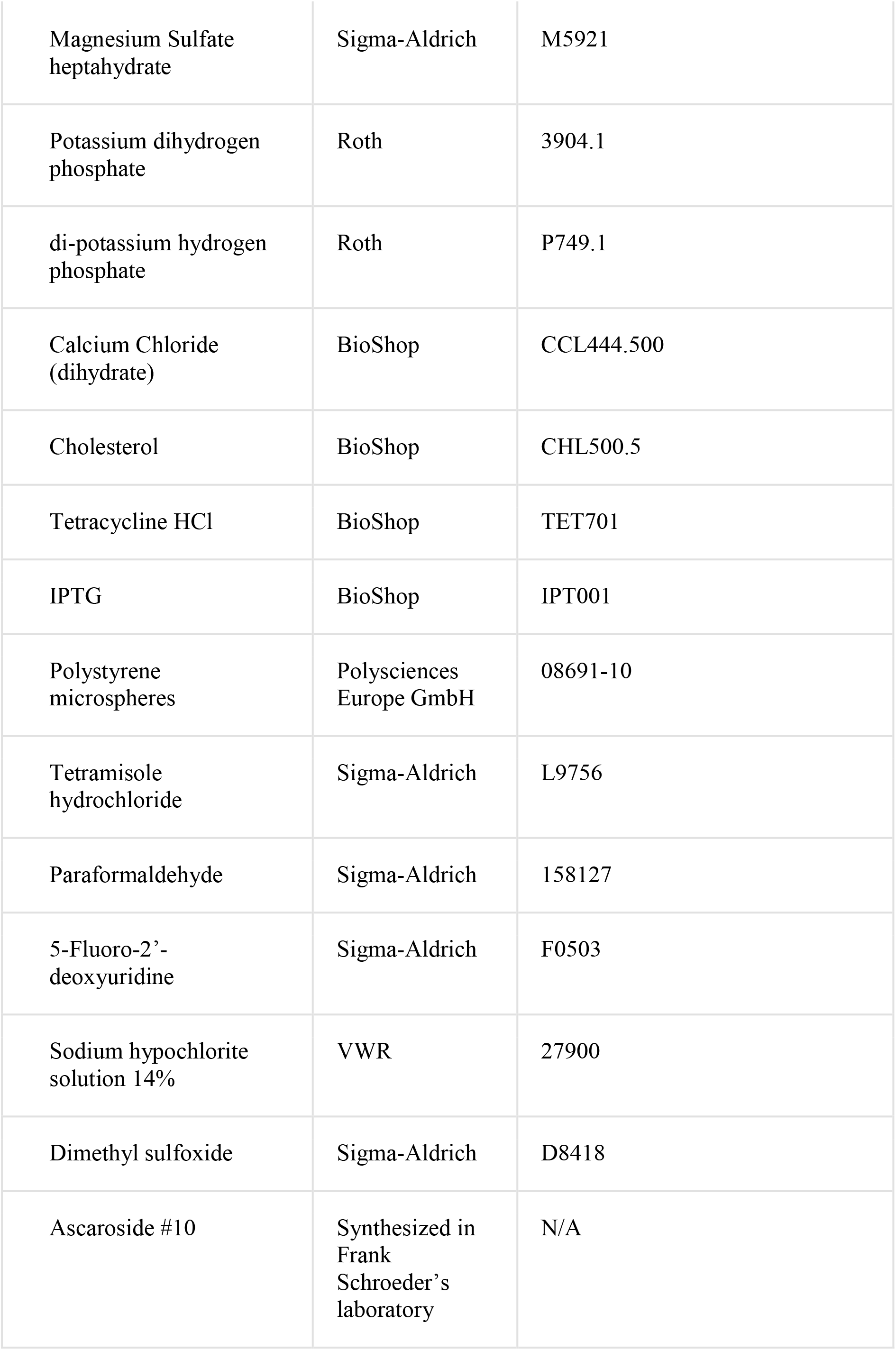
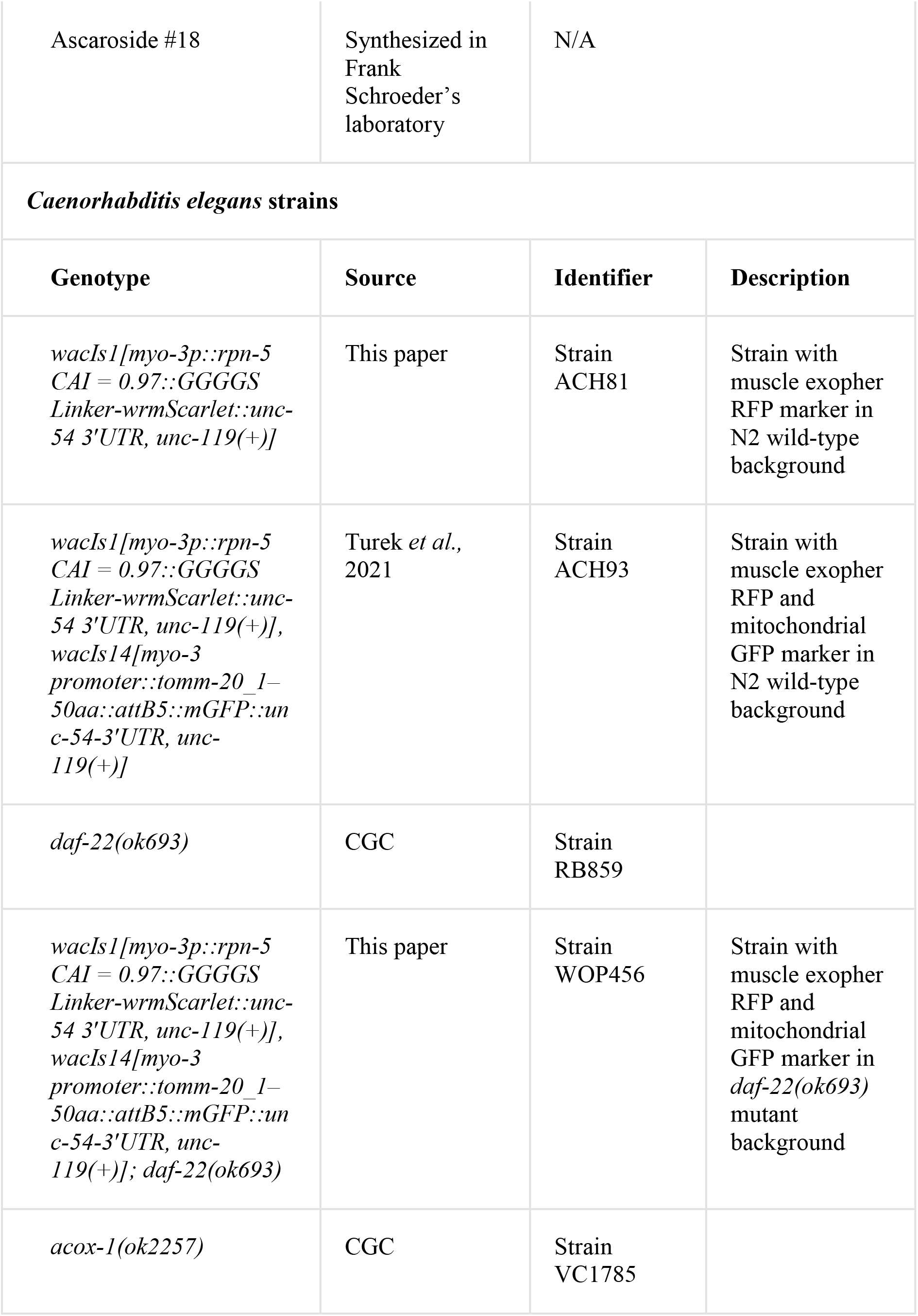
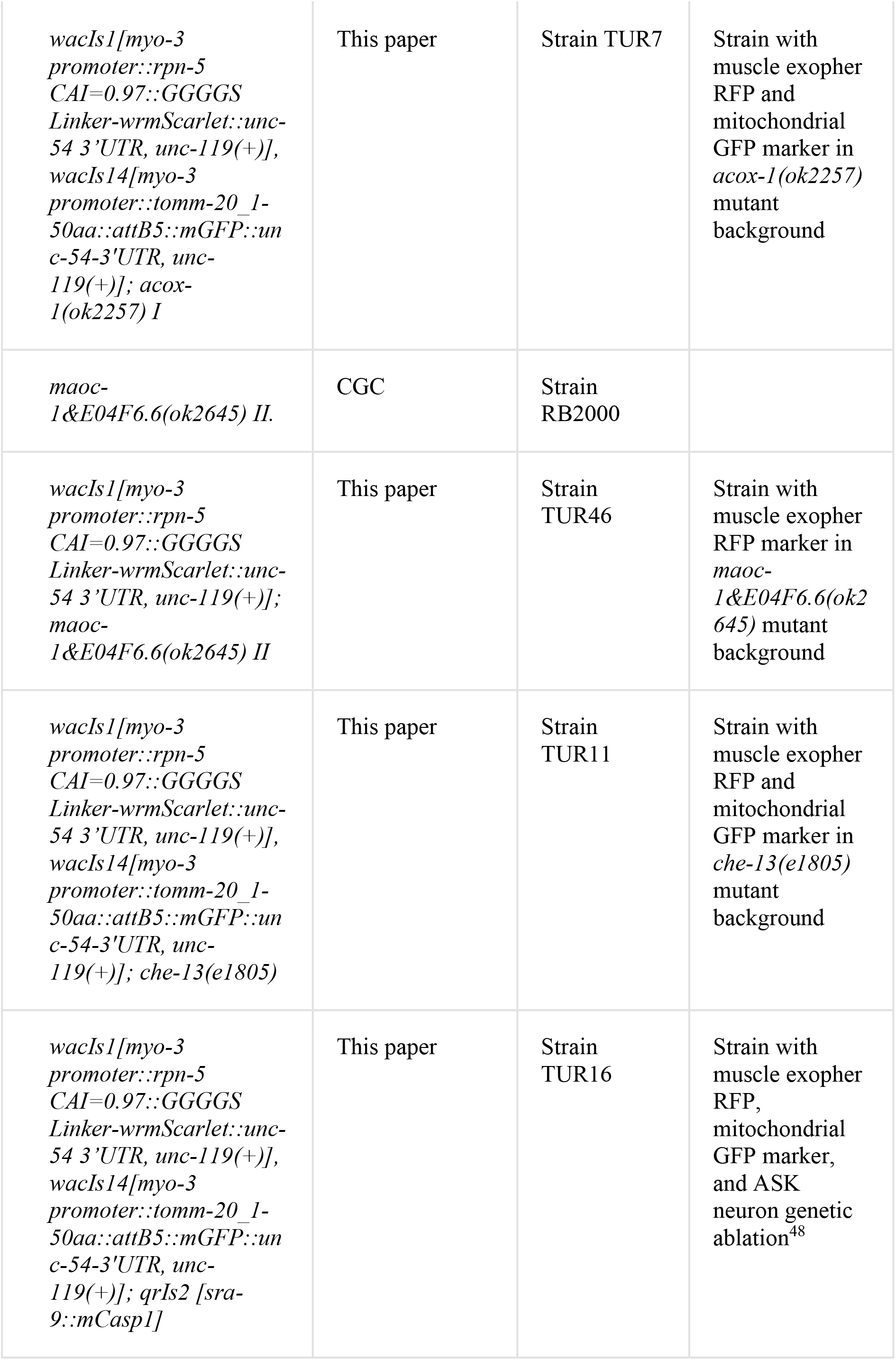
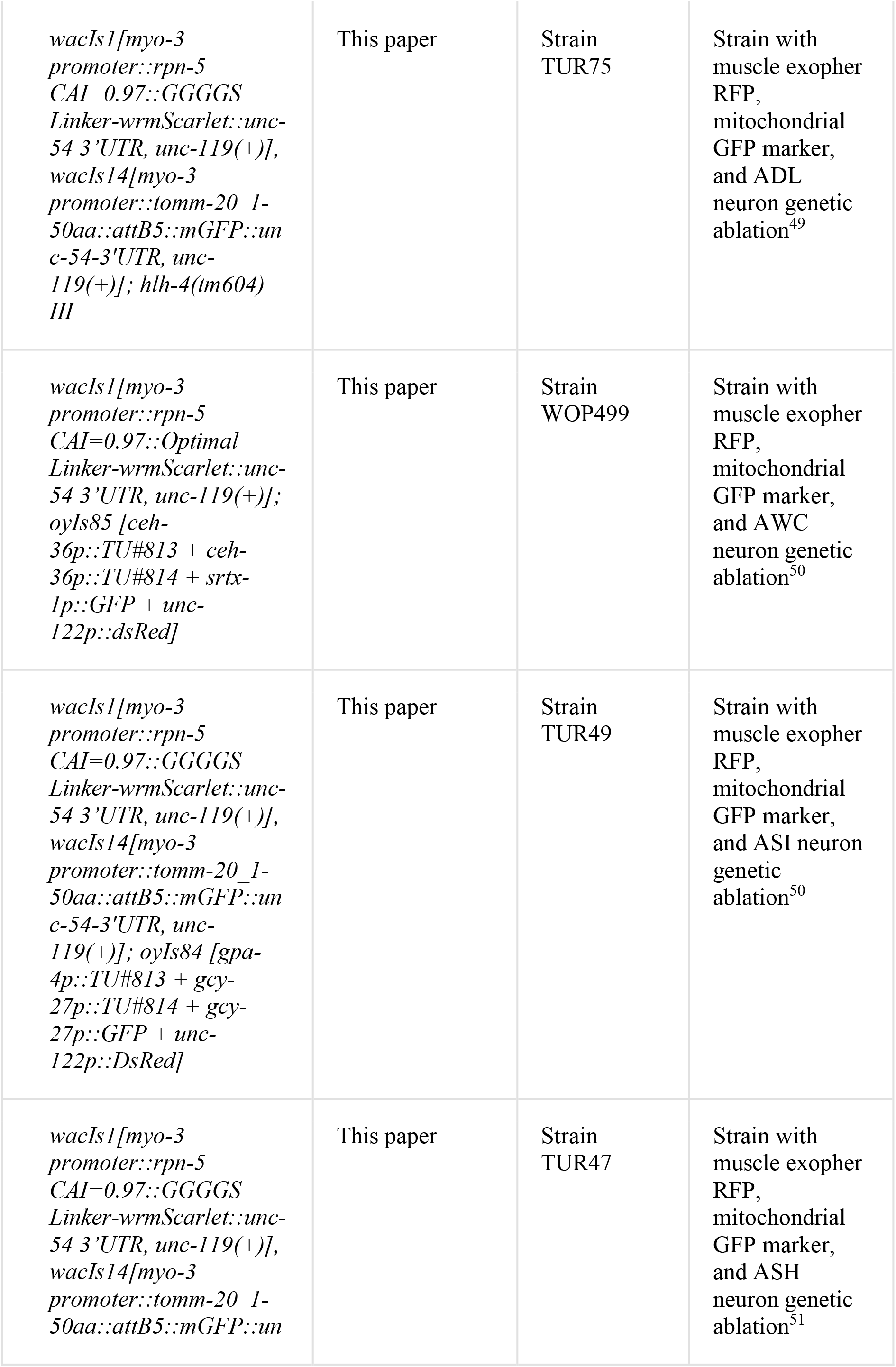
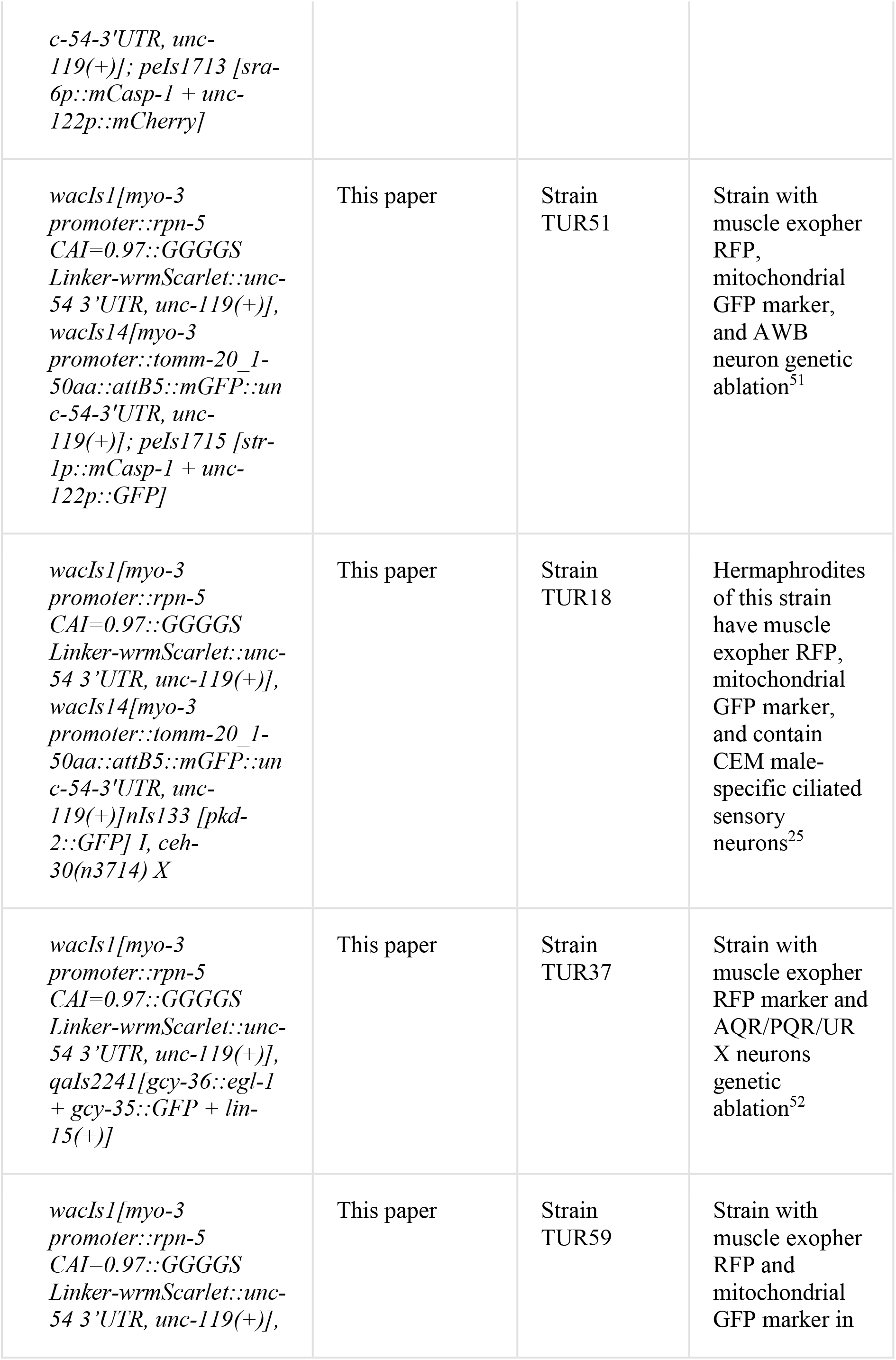
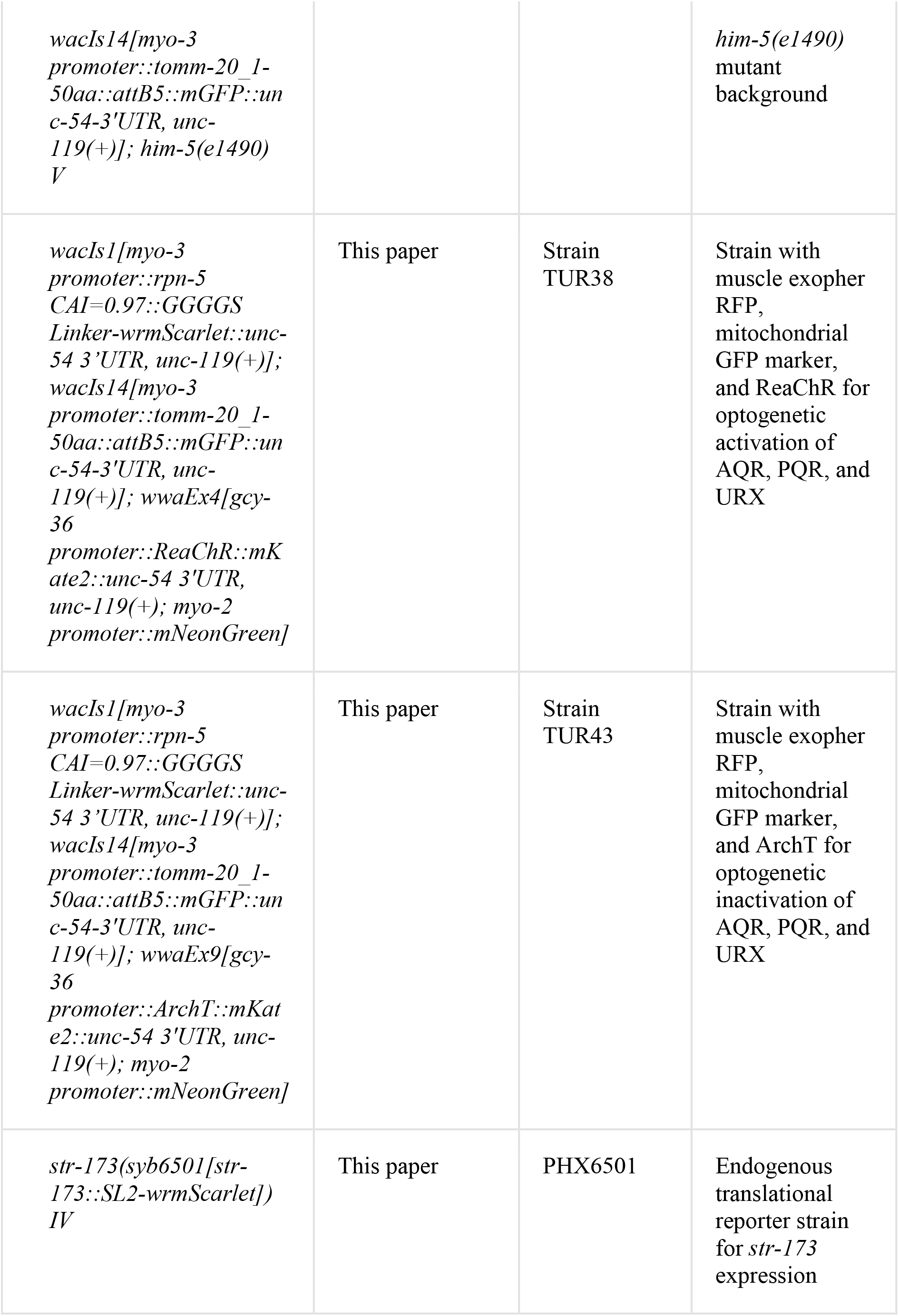
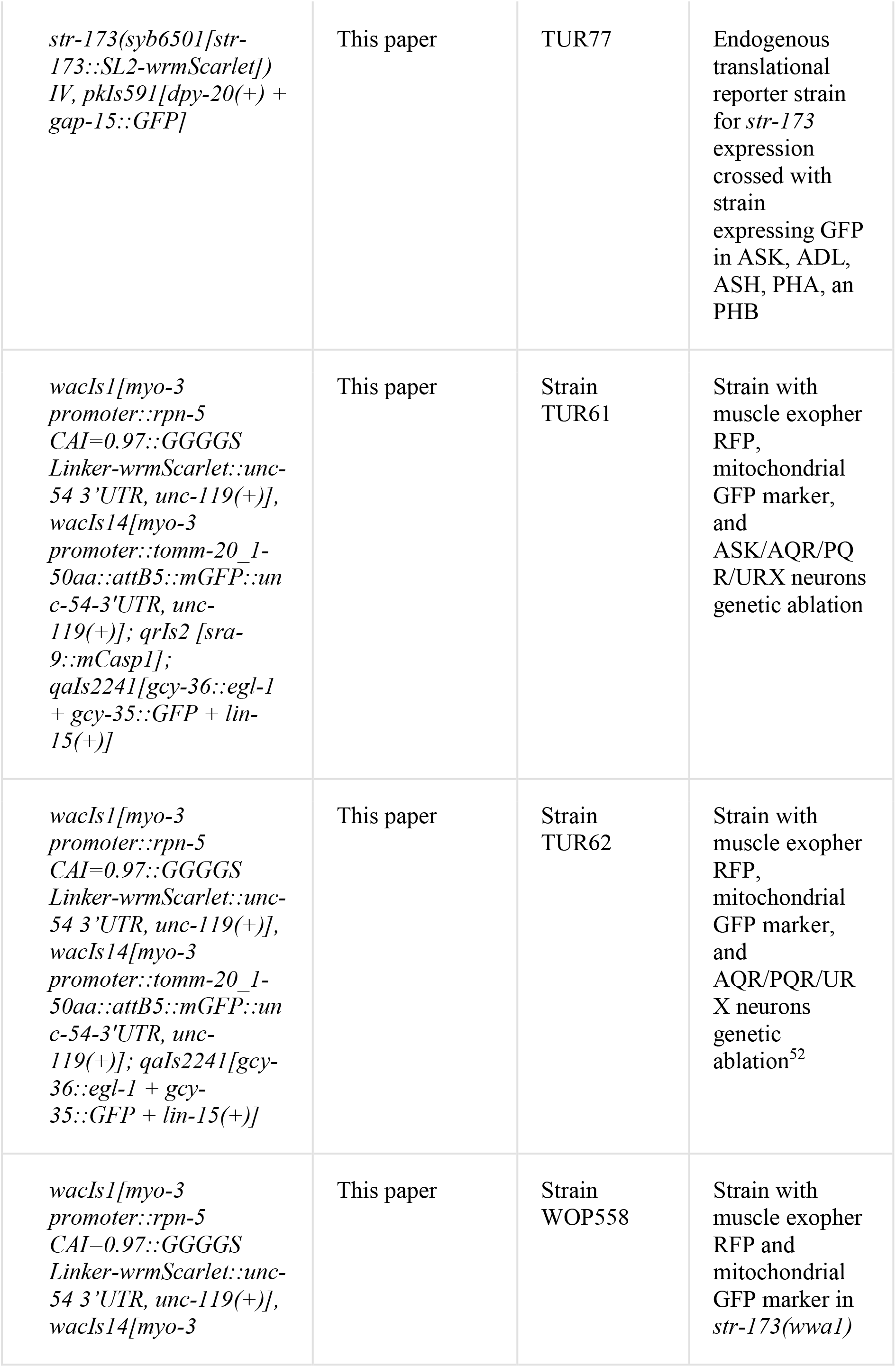
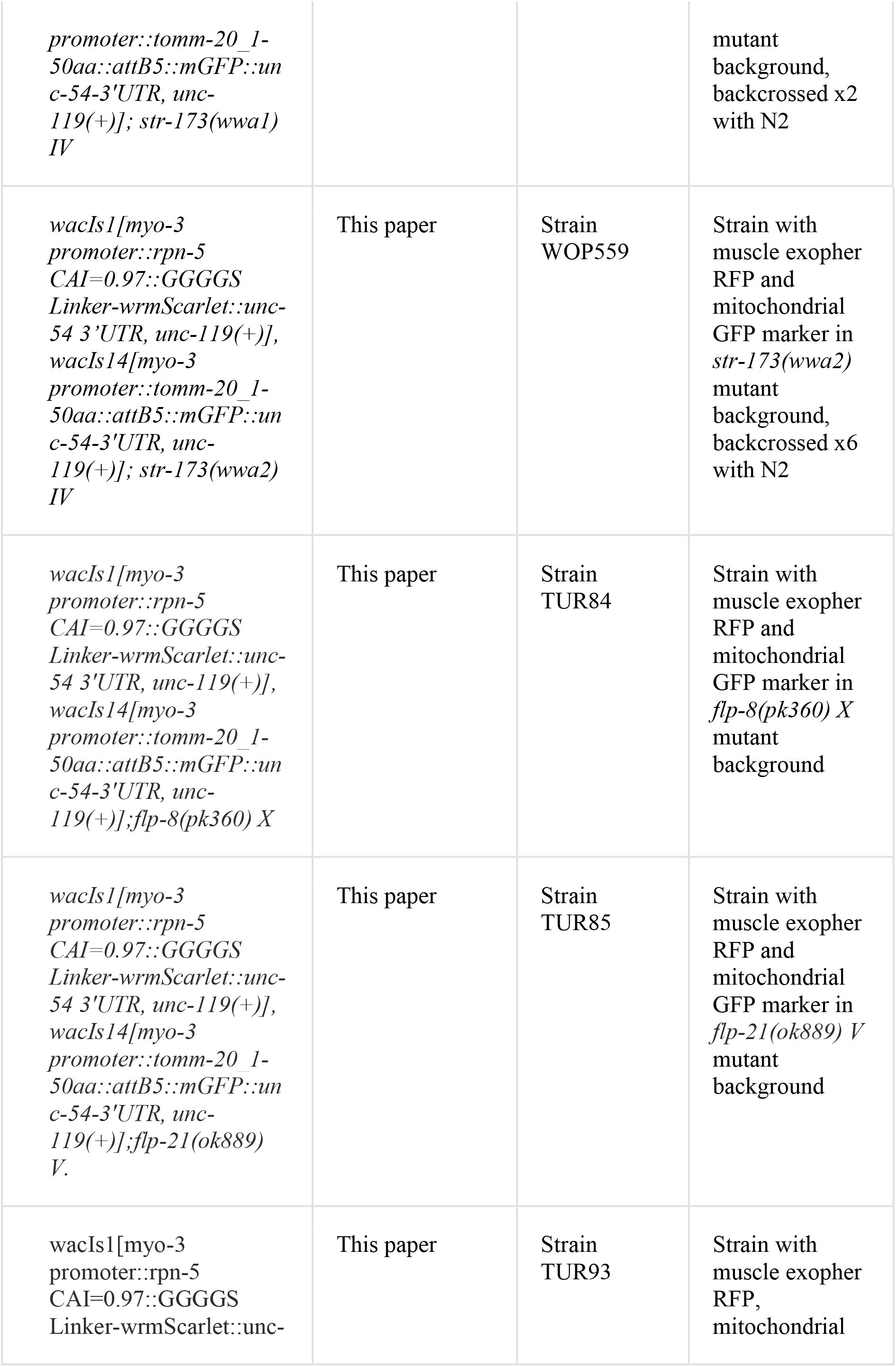
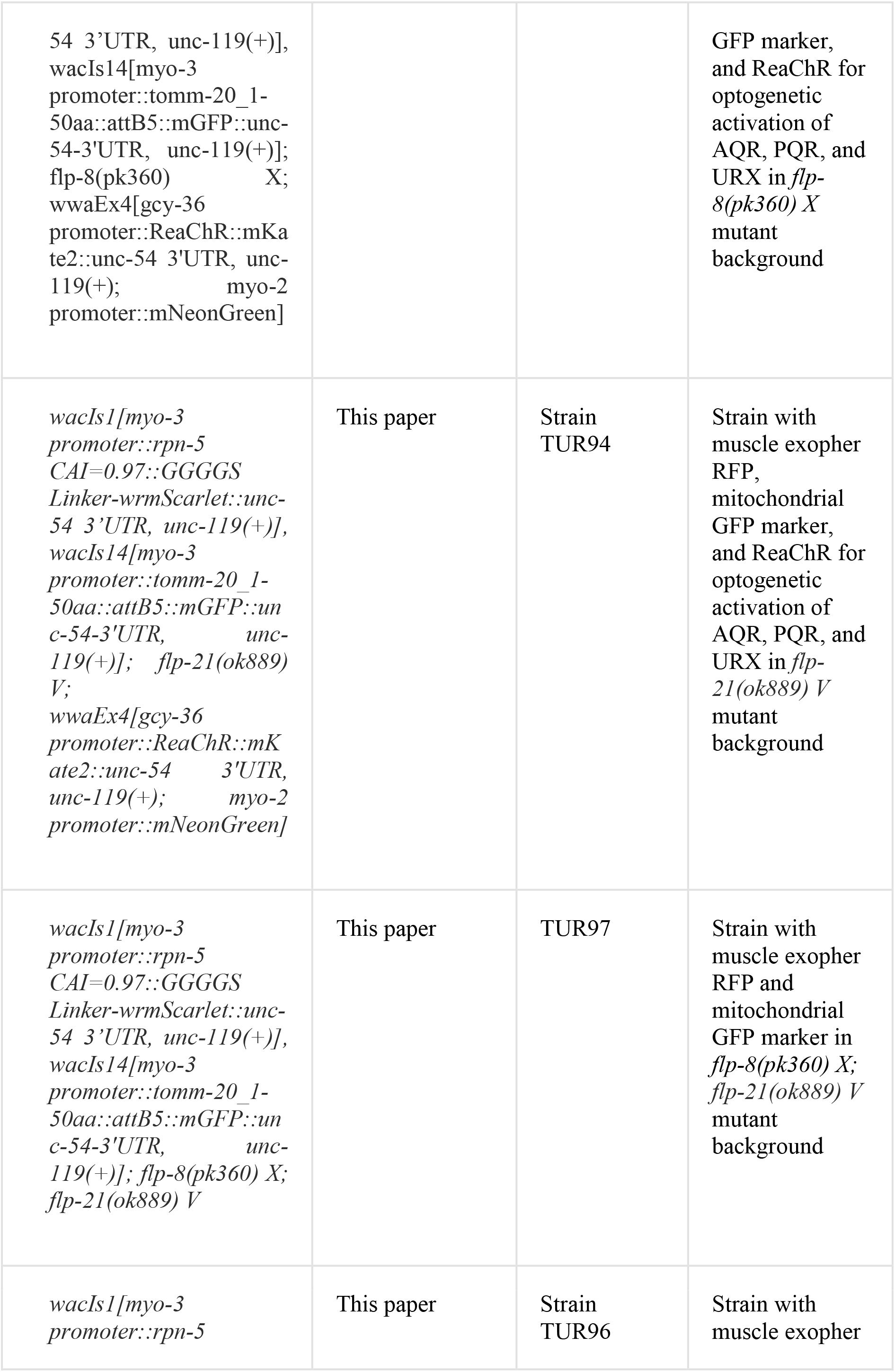
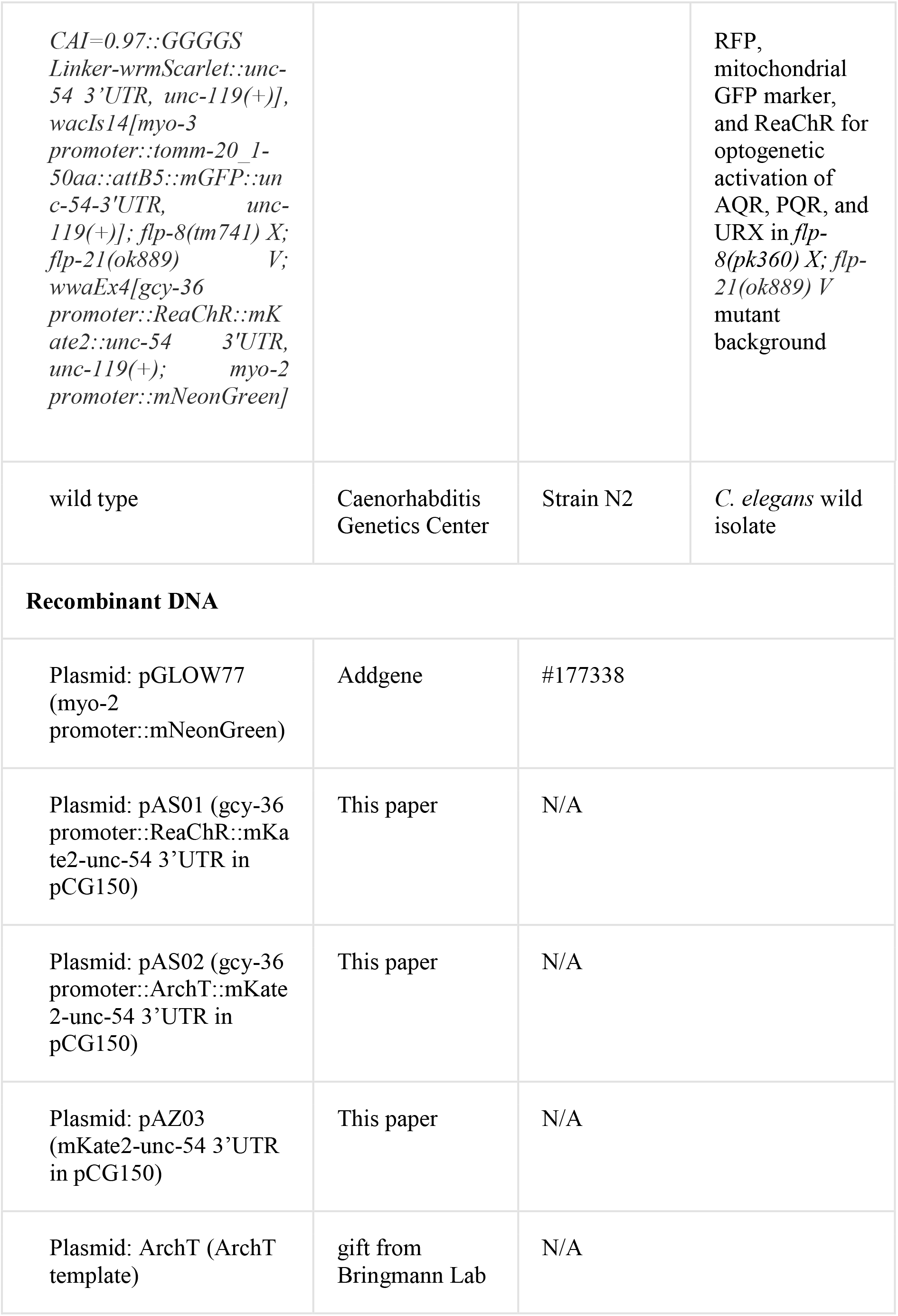
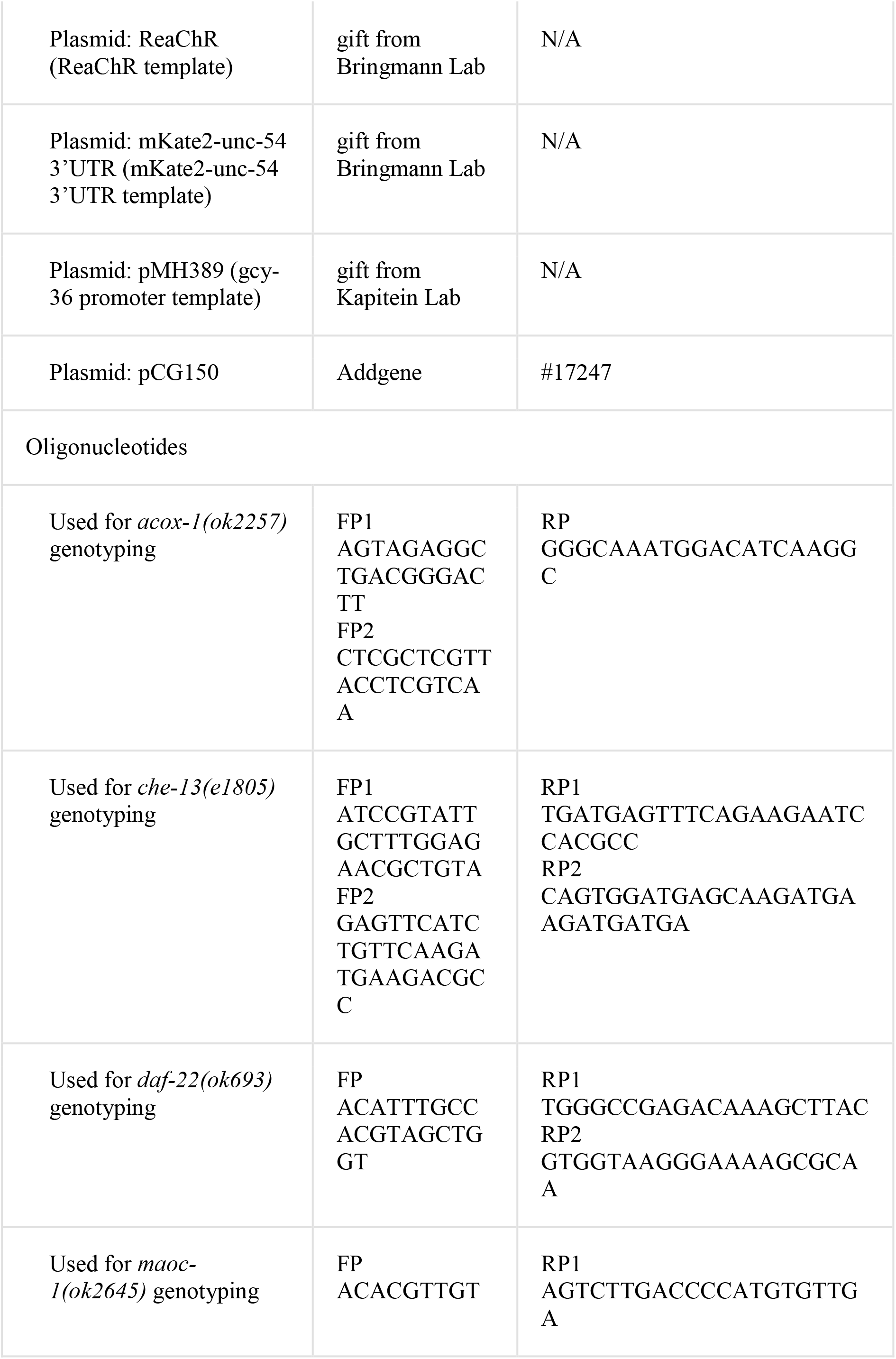
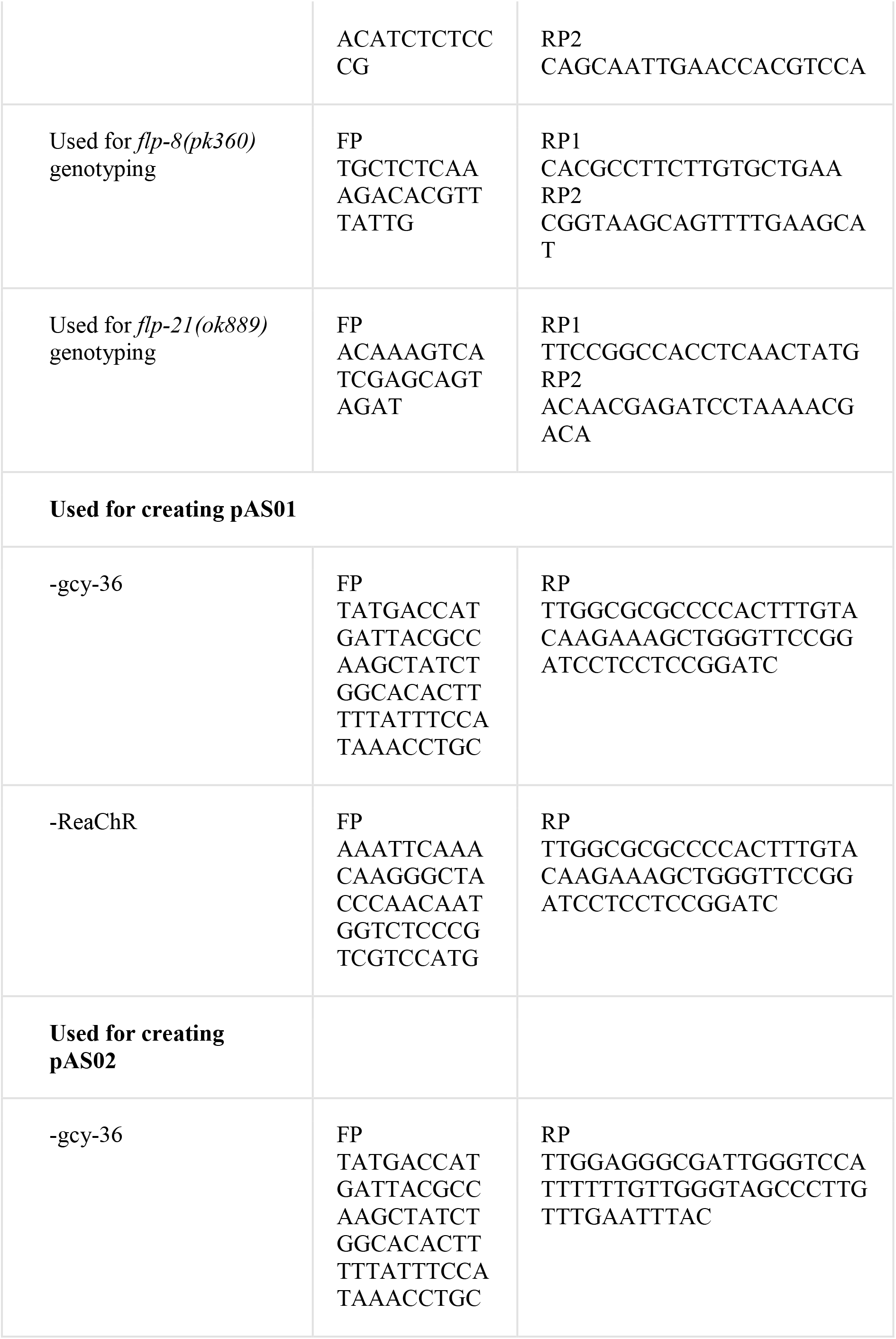
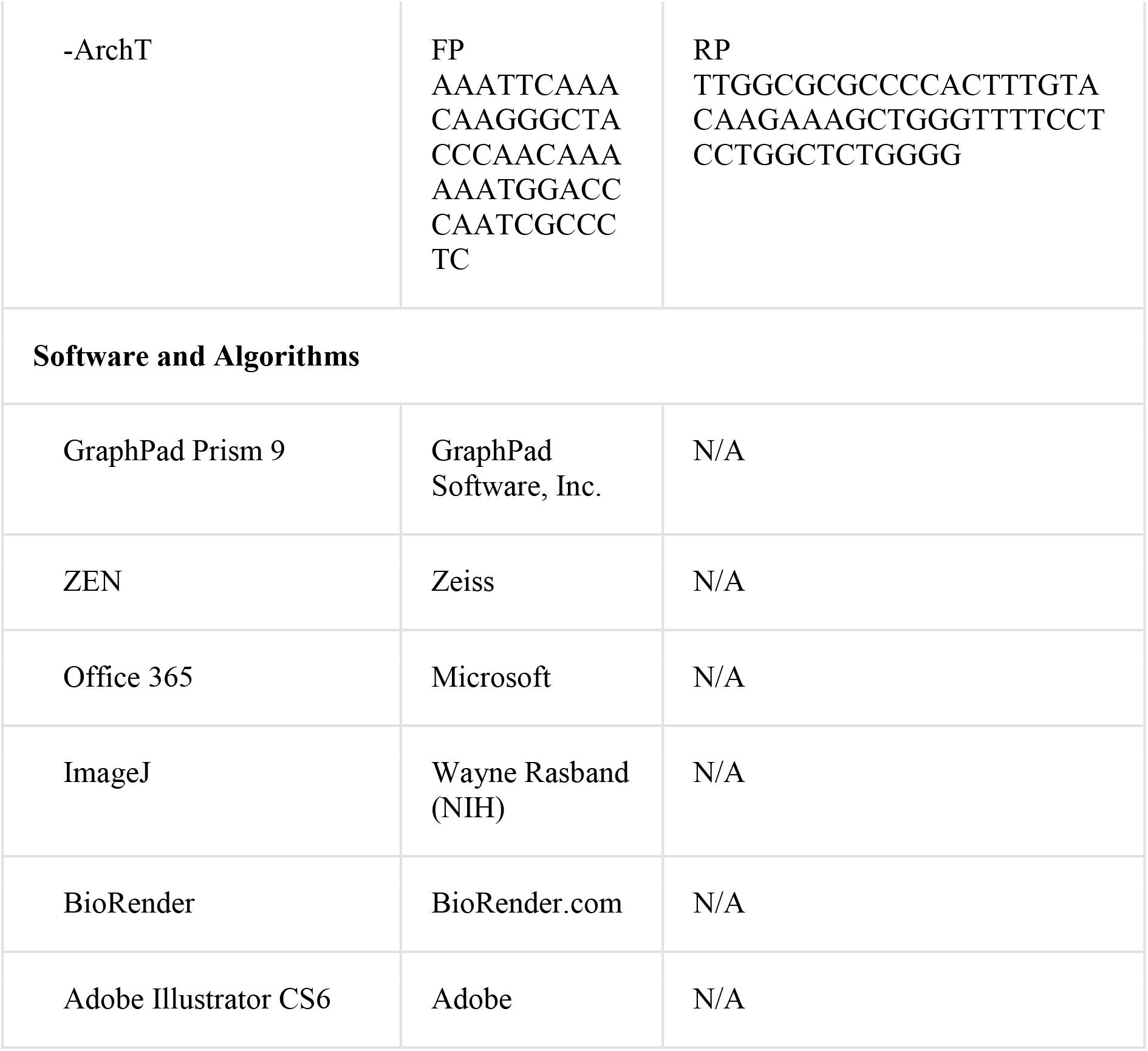

## Methods and Protocols

### Worm maintenance and strains

Worms were maintained on nematode growth medium (NGM) plates seeded with bacteria *Escherichia coli* OP50 or HT115 at 20°C unless stated otherwise^19^. A list of all *C. elegans* strains used in the study is provided in the table at the beginning of the “Materials and Methods” section.

### Scoring exophers and fluorescence microscopy

For the assessment of exophers number, the protocol described in Turek *et al*. and Banasiak *et al*. (Bio-Protocol, detailed, step-by-step guide) was applied. Briefly, exophers were scored using stereomicroscope Zeiss Axio Zoom.V16 with filter sets 63 HE and 38 HE. For each exopher scoring assay, worms were age-synchronized either from pretzel-stage embryos, or larval stages L1 and L4. On adulthood day-2, animals were visualized on NGM plates, and the number of exophers was counted in each freely moving worm.

The representative pictures presented in the manuscript were acquired with an inverted Zeiss LSM800 laser-scanning confocal microscope with a 40x oil immersion objective. To excite the GFP and RFP fluorescent proteins, 488- and 561-nm lasers were used. For visualization, animals were immobilized on 3% agarose pads with 6µl of PolySciences 0.05 µm polystyrene microspheres or 25 µM tetramisole.

### RNA interference assay

RNA interference in *C*. *elegans* was performed using the standard RNAi feeding method and RNAi clone^53^, previously described in Turek *et al.,* 2021^6^. Briefly, NGM plates, supplemented with 12,5µg/ml tetracycline, 100µg/ml ampicillin, and 1mM IPTG, were seeded with *E. coli* HT115 bacteria containing dsRNA against the gene of interest. Control group animals were fed with bacteria containing the empty vector L4440. Age-synchronized pretzel-stage embryos or L1 larvae were placed on freshly prepared plates and cultured until day 2 of adulthood when exophers were scored. The age of the worms was verified at the L4 larvae stage, either younger or older worms were removed from the experiment.

### Pre-conditioning of the plates with *C. elegans* males

50 males from the strain with *him-5(e1490)* mutation were transferred on a fresh, 35 mm NGM plate with *E. coli* OP50 or HT115 at the L4 developmental stage and older. After 48 hours, males were removed from the plates.

#### Culturing worms from L1 to adulthood day 2 on plates pre-conditioned with males

L1 larvae were transferred to plates pre-conditioned with males in a group of 10 hermaphrodites per plate. For the control group, L1 larvae were transferred to fresh, 35 mm NGM plates seeded with *E. coli* OP50 or HT115 without pre-conditioning with males. Worms were cultured up to adulthood day 2 when muscle exophers were counted in each animal.

#### Culturing worms in 24 hours intervals from L1 to adulthood day 2 on plates pre-conditioned with males

Worms were kept on plates pre-conditioned with males only for 24 hours at different developmental stages. All hermaphrodites were picked at L1 stage and grouped into 10 worms per plate population. The first group was cultured on pre-conditioned plates for 24 hours starting from hatching from the egg. After 24 hours worms were transferred to a fresh NGM plate seeded with OP50 bacteria. The second group was grown for 24 hours after hatching from the egg on the NGM plate, then the worms were moved to pre-conditioned with males plates for 24 hours, after that time they were transferred to fresh NGM plates seeded with OP50 bacteria. The third group was cultured for the first 48 hours on NGM plates, then transferred to pre-conditioned with males plates for 24 hours, and then replated on fresh NGM plates. The fourth group was cultured from the L1 stage for 72 hours on standard NGM plates, after that time worms were transferred to plates pre-conditioned with males for 24 hours. All experimental groups were cultured from L1 to adulthood day 2 when muscle exophers were counted.

### Egg retention assay

Firstly, using the stereomicroscope, muscle exophers were counted in each worm. In the following step, hermaphrodites were exposed to a 1.8% hypochlorite solution. When they dissolved, embryos retained in the uterus were counted.

### Metabolic inactivation of bacterial food source

The preparation of plates with a metabolically inactive food source for the worms to determine its effects on exophergenesis was done according to the protocol described in Beydoun *et al.*, 2021^54^. Briefly, a single *E. coli* HT115 or OP50 colony was inoculated overnight and then the bacterial culture was split into two flasks. Next, paraformaldehyde (PFA) was added to a final concentration of 0.5% in only one of them, and the flask was placed in the 37°C shaking incubator for 1 h. Afterward, the aliquots were transferred to 50 mL tubes, centrifuged at approximately 3000 × g for 20 min, and washed with 25 mL of LB five times. Control and PFA-treated bacteria were later concentrated accordingly and seeded on the NGM plates.

As control of PFA treatment, new bacterial cultures in LB were set in the 37°C shaking incubator overnight to ensure the replication was blocked and there was no bacterial growth.

Plates were used for experiments at least 5 days after their preparation to ensure no bacteria-derived metabolites were left on plates with PFA-treated *E. coli*. A new batch of bacterial food source was prepared for each biological repetition.

### Brood size quantification

Age-synchronized hermaphrodites at the L4 developmental stage were picked for the experiment. Worms were cultured as single worms per 60 mm NGM plate, seeded with *E. coli* OP50 or HT115. Ten worms were counted as one biological repeat. Worms were transferred to fresh plates every day from adulthood day 1 until the end of the egg laying period.

The number of eggs laid over the worms’ reproductive lifetime was counted manually daily. The data is presented as the total number of eggs laid by each animal.

### Quantifying the number of exophers in worms grown in different temperatures

Worms were confronted with a range of physiological temperatures: low 15 °C, optimal 20 °C, and high 25 °C throughout their development until exophers were scored. To compare the corresponding stages of development at various temperatures, worms were additionally sorted at the L4 stage. The exopher number was assessed based on the timepoint of maximal egg laying, approx. 140 h or 78 h from egg hatching at low or high maintenance temperatures, respectively. Timing calculations were based on the *C. elegans* development timeline at different temperatures^55^.

### Culturing worms in different population sizes

L1 larvae were transferred to 35 mm NGM Petri dishes seeded with *E. coli* OP50 or HT115 bacteria strains (100 μL of bacteria from 10 mL overnight culture grown in 50 mL Erlenmeyer flask) immediately after hatching as a single larva or in a group of 5, 10 or 100 animals. In total, thirty plates with single worms, six plates with 5 worms, three plates with 10 worms, and one plate with 100 worms were used per biological repeat. Exophers were scored on adulthood day 2 in the worms expressing RFP in BWM.

### Quantifying the influence of molecules released by ascaroside biosynthesis mutant on exopher production in wild-type worms

Nine freshly hatched L1 larvae from selected ascaroside biosynthesis mutant strains (*maoc-1(ok2645), daf-22(ok693),* or *acox-1(ok2257)*) or wild-type hermaphrodites (as a control) were transferred to fresh NGM plates. Next, one L1 larva of a reporter strain expressing RFP in BWM was added to each plate and 10 worms in total were grown together on 35 mm Petri dish seeded with *E. coli* OP50 or HT115 bacteria strains. On adulthood day 2 the number of exophers was quantified in the worms expressing RFP in BWM, according to description.

### Generation of *str-173* mutant strains

The *str-173* gene mutants (*str-173(wwa1)* and *str-173(wwa2)*) were generated using CRISPR/Cas9 method as previously described^56^. The crRNA sequence used was ATAATTGGTGGATATACAAATGG. The *str-173* gene locus was sequenced, and deletions were mapped to the first exon (Extended Data Figure 3d). Both mutations cause frame shifts and are most likely molecular null alleles. Strains were backcrossed twice (strain *str-173(wwa1)* and 6 times *(str-173(wwa2)*) against N2 wild-type worms.

### Generation of optogenetic strains

Optogenetic strains created for this paper contain red-shifted Channel Rhodopsin (ReaChR) or archaerhodopsin from Halorubrum strain TP009 (ArchT). To generate these strains, firstly, mKate2-unc-54 3’UTR was amplified from the template and cloned into pCG150 to create a pAZ03 plasmid. Next, the *gcy-36* promoter was amplified from pMH389, and ReaChR and ArchT were amplified from respective templates. The *gcy-36* promoter was then cloned into pAZ03 plasmid with ReaChR and ArchT separately. As a result two plasmids were created: gcy-36 promoter::ReaChR::mKate2-unc-54 3’UTR in pCG150 and gcy-36 promoter::ArchT::mKate2-unc-54 3’UTR in pCG150. Plasmids were sequenced to verify the correct sequence of the cloned constructs. All constructs generated for this study were made using the SLiCE method^57^.

Transgenic strains with extrachromosomal arrays were generated by microinjection. DNA was injected into exopher reporter strain worms with muscle exopher RFP and mitochondrial GFP marker (ACH93). For injection, DNA was prepared as follows: construct 90 ng/µL and co-injection marker 10 ng/µL. Positive transformants were selected according to the presence of co-injection markers (myo-2 promoter::mNeonGreen).

The constructs created for this project and primers used for amplification are listed in the table at the beginning of the “Materials and Methods” section.

### Optogenetics assays

#### On adulthood day 2

For optogenetic activation or inhibition, 35-mm NGM plates seeded with HT115 *E. coli* bacteria were covered with 0.2 µM all-trans retinal (ATR). Control plates were not covered with ATR. Ten to twelve age-synchronized worms were picked per plate from optogenetic strains (expressing ReaChR or ArchT in AQR/PQR/URX neurons) at adulthood day 1. After 24-hour incubation at 20°C and darkness, muscle exophers extruded by worms were counted **(1)**. Next, experimental plates were placed on the stereomicroscope and illuminated for 1 hour with green light (HXP 200C illuminator as a light source, band-pass filter Zeiss BP 572/25 (HE), the green light intensity measured at 561 nm = 0.07 mW/mm^2^). Immediately after illumination muscle exophers were counted **(2)**. Subsequently, exophers scorings were performed 15 min **(3)** and 30 min **(4)** after the illumination with a green light was completed. Worms were kept in the darkness in between counts. The control group was not illuminated. To provide similar environmental conditions control plates were placed next to the experimental plate but were shielded from light. Control and treated groups were randomized before the start of the experiment.

#### On adulthood day 1

The assay was conducted similarly as on adulthood day 2 with a difference in time of exposure to the light. Twelve age-synchronized worms from optogenetic strains were picked at the L4 developmental stage and transferred on a 35-mm NGM plate covered with ATR (no ATR for the control group). Worms were then incubated for 24 hours at 20 °C and darkness. Experimental plates were placed on the stereomicroscope and illuminated for 1 hour with green light (HXP 200C illuminator as a light source, band-pass filter Zeiss BP 572/25 (HE), the green light intensity measured at 561 nm = 0.07 mW/mm^2^). Exophers extruded from BWM were counted before the exposure to the light **(1)**, immediately after the exposure to the light **(2)**, and 15 min **(3)**, 30 min **(4),** and 24 hours **(5)** after the exposure to the light.

### FUdR assay

Age-synchronized young adult animals (day 0) were placed on NGM plates containing 25 μM fluorodeoxyuridine (FUdR) or control NGM plates without FUdR. Exophers number were scored when worms reached adulthood day 2 using a stereomicroscope.

### Transcriptome analysis

RNA extractions, library preparations, and sequencing were conducted at Azenta US, Inc (South Plainfield, NJ, USA) as follows:

#### RNA Extraction

Total RNA was extracted using Qiagen RNeasy Plus mini kit following the manufacturer’s instructions (Qiagen, Hilden, Germany).

#### Library Preparation with polyA selection and Illumina Sequencing

Extracted RNA samples were quantified using Qubit 2.0 Fluorometer (Life Technologies, Carlsbad, CA, USA) and RNA integrity was checked using Agilent TapeStation 4200 (Agilent Technologies, Palo Alto, CA, USA).

RNA sequencing libraries were prepared using the NEBNext Ultra II RNA Library Prep Kit for Illumina following the manufacturer’s instructions (NEB, Ipswich, MA, USA). Briefly, mRNAs were first enriched with Oligo(dT) beads. Enriched mRNAs were fragmented for 15 minutes at 94 °C. First strand and second strand cDNAs were subsequently synthesized. cDNA fragments were end repaired and adenylated at 3’ends, and universal adapters were ligated to cDNA fragments, followed by index addition and library enrichment by limited-cycle PCR. The sequencing libraries were validated on the Agilent TapeStation (Agilent Technologies, Palo Alto, CA, USA), and quantified by using Qubit 2.0 Fluorometer (Invitrogen, Carlsbad, CA) as well as by quantitative PCR (KAPA Biosystems, Wilmington, MA, USA).

The sequencing libraries were multiplexed and loaded on the flowcell on the Illumina NovaSeq 6000 instrument according to the manufacturer’s instructions. The samples were sequenced using a 2×150 Pair-End (PE) configuration v1.5. Image analysis and base calling were conducted by the NovaSeq Control Software v1.7 on the NovaSeq instrument. Raw sequence data (.bcl files) generated from Illumina NovaSeq was converted into fastq files and de-multiplexed using Illumina bcl2fastq program version 2.20. One mismatch was allowed for index sequence identification.

#### Sequencing Data Analysis

After investigating the quality of the raw data, sequence reads were trimmed to remove possible adapter sequences and nucleotides with poor quality using Trimmomatic v.0.36. The trimmed reads were mapped to the *Caenorhabditis elegans* reference genome available on ENSEMBL using the STAR aligner v.2.5.2b. The STAR aligner is a splice aligner that detects splice junctions and incorporates them to help align the entire read sequences. BAM files were generated as a result of this step. Unique gene hit counts were calculated by using feature Counts from the Subread package v.1.5.2. Only unique reads that fell within exon regions were counted.

After the extraction of gene hit counts, the gene hit counts table was used for downstream differential expression analysis. Using DESeq2, a comparison of gene expression between the groups of samples was performed. The Wald test was used to generate p-values and Log2 fold changes. Genes with adjusted p-values < 0.05 and absolute log2 fold changes > 1 were called as differentially expressed genes for each comparison.

RNAseq data was deposited in the GEO database (GSE241786) and can be accessed using the following links: https://www.ncbi.nlm.nih.gov/geo/query/acc.cgi?acc=GSE241786

### Growing hermaphrodite on plates pre-conditioned with other hermaphrodites

50 wild-type hermaphrodites at the L4 stage were placed on a fresh, 35 mm NGM plate seeded with *E. coli* OP50 bacteria (100 μL of bacteria from 10 mL overnight culture grown in 50 mL Erlenmeyer flask). After 12-16 hours, hermaphrodites were removed from the plate and a single pretzel-stage embryo was transferred to the hermaphrodite-conditioned plate. For the control groups single pretzel-stage embryos or ten pretzel-stage embryos were situated on a fresh, 35 mm NGM plate with *E. coli* OP50 bacteria without pre-conditioning with hermaphrodites. When worms reached the second day of adulthood, the number of exophers was scored in each animal.

### Quantifying the influence of larval pheromones on exopher production in hermaphrodites

A single pretzel-stage embryo was placed on a fresh, 35 mm NGM plate seeded with *E. coli* OP50 bacteria (100 μL of bacteria from 10 mL overnight culture grown in a 50 mL Erlenmeyer flask). The worm was cultured until the late L4 stage, when 50 L1 larvae were added to a single worm. For the control groups, single pretzel-stage embryos or ten pretzel-stage embryos were placed on a fresh, 35 mm NGM plate with *E. coli* OP50 bacteria. Muscle exophers were scored in each animal at adulthood day 2.

### Quantifying the influence of molecules released by ascaroside biosynthesis mutant on exopher production

Nine freshly hatched L1 larvae from selected ascaroside biosynthesis mutant strains (*maoc-1*(ok2645), *daf-22*(ok693), or *acox-1*(ok2257)) or wild-type hermaphrodites (as a control) were transferred to fresh NGM plates. Next, one L1 larva of a reporter strain expressing RFP in body wall muscles was added to each plate, and 10 worms in total were grown together up to adulthood day 2. Muscle exophers were counted in each animal expressing RFP in BWM.

### Growing hermaphrodites on plates supplemented with synthetic ascr#10 or ascr#18

Synthetic, concentrated ascr#10 and ascr#18 were stored in DMSO at -80°C. The stocks were diluted to working solutions with water. A total of 100µL (containing 1ng or 1pg) was applied and rubbed into the 35-mm NGM plate with a sterile glass rod. Plates were incubated at 20 °C overnight. The following day plates were seeded with 100µL of *E.coli* OP50 bacteria, and incubated for an additional 24 hours at 20 °C. Control plates were prepared similarly using water instead of an ascaroside solution. Pretzel-stage eggs were transferred on plates supplemented with ascarosides and control plates in groups of twelve eggs per plate. On day 2 adulthood the number of exophers was quantified in the worms expressing RFP in BWM.

### Data analysis and visualization tools

The exophers were scored at adulthood day 2 using a stereomicroscope unless stated otherwise. Data analysis was performed using Microsoft® Excel® and GraphPad Prism 9 software. Graphical representation of data was depicted using GraphPad Prism 9.

### Statistical analysis

No statistical methods were used to predetermine the sample size. Worms were randomly allocated to the experimental groups for all the data sets and experiments were performed blinded for the data sets presented in the following figures: Fig. 1; Fig. 3c, d, e, f, g, h; Fig. 4a, d, g; Fig. 5c, d, e, f, g, h, i, j; Fig. 7a, b; Supplementary Fig. 3a, b. Non-Gaussian distribution of residuals was assumed; therefore, nonparametric statistical tests were applied: Mann– Whitney (in comparison between two groups) or Kruskal-Wallis test with Dunn’s multiple comparisons test (in comparison between more than two groups). *P*-value < 0.05 is considered significant.

## Data availability statement

The data that support the findings of this study are available from the corresponding authors upon reasonable request. RNAseq data was deposited in the GEO database (GSE241786) and can be accessed using the following links: https://www.ncbi.nlm.nih.gov/geo/query/acc.cgi?acc=GSE241786

## Acknowledgements

Some strains were provided by the CGC, which is funded by NIH Office of Research Infrastructure Programs (P40 OD010440). We thank Henrik Bringmann and Lukas Kapitein for plasmids; Peter Askjaer, Henrik Bringmann, and Antonio Miranda Vizuete for discussions and comments on the manuscript; Zofia Olszewska, Marta Niklewicz, and Monika Woźniak for assistance with worms maintenance. Work in the M. T. Laboratory was mainly funded by a National Science Centre SONATA grant (2019/35/D/NZ3/04091) and additionally supported by a National Science Centre SONATA BIS grant (2021/42/E/NZ3/00358). Work in the W. P. Laboratory was funded by the Foundation for Polish Science co-financed by the European Union under the European Regional Development Fund (grant POIR.04.04.00-00-5EAB/18-00 to K.O., and W.P.), and additionally supported by the European Molecular Biology Organization (EMBO Installation Grant No. 3916 to K.O., and W.P.), and the Norwegian Financial Mechanism 2014-2021 operated by the Polish National Science Centre, Poland (project contract number 2019/34/H/NZ3/00691 to W.P.).

## Author contributions

Conceptualization: MT, WP; Data curation: MT, WP; Formal analysis: AS, KO, KK, MT, WP, FCS, JY, AIT, LA; Funding acquisition: MT, WP; Investigation: AS, KO, KK, YI, AIT, LA, MT; Methodology: MT, WP, KO, AS, KK, FCS, YI; Project administration: MT, WP; Resources: MT, WP; Supervision: MT, WP; Validation: MT, WP, FCS, AS, KO, KK; Visualization: AS, KO, KK, MT; Writing – original draft: MT, WP; Writing – review & editing: MT, WP, FCS, AS, KO, KK.

